# High-speed 3D single-virus tracking reveals actin-aided viral trafficking of SARS-CoV-2 on the plasma membrane

**DOI:** 10.64898/2026.04.03.716319

**Authors:** Yuxin Lin, Xinrui Lu, Jack Exell, Haoting Lin, Courtney Johnson, Kevin D. Welsher

## Abstract

Early interactions between viruses and live cells are difficult to resolve due to rapid extracellular motion, 3D nature of the cell membrane, and the fast, nanoscale interactions involved. While actin is a central regulator of viral entry, direct observations of actin-aided trafficking have been restricted to membrane protrusions on glass surfaces given the limitations of conventional methods. Here, high-speed 3D Tracking and Imaging microscopy (3D-TrIm) is integrated with highly photostable StayGold-labeled SARS-CoV-2 virus-like particles to capture long-term, high-resolution single-virus trajectories in live cells. This approach revealed distinct regimes of viral dynamics, including extracellular diffusion, protrusion-based surfing, and an unreported linear trafficking mode along the plasma membrane that precedes viral internalization. This work demonstrates that this membrane trafficking is actin-driven and positively correlated with ACE2 expression. These findings reveal new actin exploitation by viruses and demonstrate the utility of 3D-TrIm for dissecting dynamic virus–cell interactions at high spatiotemporal resolution.

## Introduction

Actin is a family of multifunctional proteins present in most eukaryotic cells, playing essential roles in a wide range of cellular processes, including cell motility and division, membrane protein expression and presentation, formation and movement of endocytic/phagocytic vesicles, and virus entry and egress.^1–6^ In eukaryotic cells, actin exists in two primary forms: monomeric globular G-actin and polymerized filamentous F-actin, between which rapid polymerization and depolymerization occur. The polymerization dynamics are regulated by the small GTPase RhoA and actin-interacting proteins, such as formins^7^ and actin-related protein 2/3 (Arp2/3) complexes.^8^ Within cells, actin is organized into distinct actin-rich structures that support various biological functions, including the actin cortex, microvilli, stress fibers, and protrusive structures such as filopodia.^9–12^

Given its versatility, it is not surprising that viruses actively hijack and manipulate the actin network during infection. For example, viruses have been observed to rapidly “surf” along filopodia to approach the cell body, a mechanism that facilitates subsequent viral entry.^4, 13–15^ These actin-based viral trafficking events have been reported on actin-rich cellular protrusions, especially filopodia, across a diverse range of viruses, including severe acute respiratory syndrome coronavirus 2 (SARS-CoV-2).^4, 16–18^ In theory, other non-protrusive actin-rich structures, such as the actin cortex and stress fibers, should also be capable of supporting similar trafficking mechanisms to aid viral transport and internalization. However, direct evidence and systematic observations of such processes remain scarce.^19^ One of the main obstacles is that those actin-rich structures stretch along the three-dimensional contours of the cell body. In contrast to the relatively simple, one- or two-dimensional nature of membrane protrusions, high spatiotemporal resolution in 3D is required to capture viral dynamics on cellular membranes. This challenge necessitates advanced imaging techniques capable of capturing rapid, nanoscale viral motion in 3D.

Single-virus tracking (SVT) is a powerful technique for studying virus–cell interactions, providing insights into viral dynamics at the single-particle level that are inaccessible through ensemble measurements.^20, 21^ In a conventional SVT workflow, fluorescently labeled viruses are excited with wide-field illumination and imaged onto a camera, and trajectories are reconstructed based on the localization of viral particles across consecutive frames.^22^ However, the relatively slow frame rate of cameras often limits the ability to capture highly dynamic processes. Moreover, most approaches, such as spinning disk confocal microscopy and light sheet microscopy, rely on sequential z-stack acquisition to achieve 3D tracking, significantly compromising temporal resolution. Therefore, conventional SVT methodologies often fall short in capturing sufficient spatiotemporal information, particularly for highly dynamic, three-dimensional processes, including the potential existence of viral trafficking on the actin cortex.

Active-feedback SVT methods overcome this limitation by actively re-centering the moving object within the detection volume in real time rather than passively acquiring frames.^23, 24^ This approach decouples localization from frame-based imaging, dramatically improving temporal resolution down to the millisecond scale, especially in 3D. One successful example of utilizing active-feedback SVT in understanding viral infection is 3D Tracking and Imaging microscopy (3D-TrIm),^25^ which combines active-feedback single-virus tracking with volumetric imaging of the cellular environment. In this study, 3D-TrIm microscopy is integrated with StayGold-labelled virus-like particles (StayGold-VLPs) to enable long-term, high-resolution tracking of single viruses. This platform allows visualization and dissection of the infection process of SARS-CoV-2 VLPs with a kilohertz sampling rate of viral position along with full volumetric images of the cellular environment. Recorded viral trajectories exhibited extracellular diffusion, surfing on cellular protrusions, and linear trafficking of internalized VLPs. In addition, for the first time, linear trafficking modes on the cell membrane were observed for single SARS-CoV-2 VLPs. This viral linear membrane trafficking was found to be actin-dependent, microtubule-independent, and correlated with receptor expression levels. These findings, only revealable by long-term high-speed single-virus tracking microscopy, expand the paradigm of viral “surfing” beyond filopodia to a more general surface transport mechanism.

## Results and Discussion

### SVT with long duration and high spatiotemporal resolution via 3D-TrIm and CoV2/SG-VLPs

The 3D-TrIm microscope is comprised of two key components (Figure 1), one which tracks the virus (3D Single-Molecule Active-feedback Real-time Tracking, 3D-SMART), and one that images the cellular context (3D Fast Acquisition by z-Translating Raster, 3D-FASTR). The implementation of 3D-SMART in the current work is similar to previous reports.^25–27^ Briefly, a pair of electro-optic deflectors (EODs) and a tunable acoustic gradient (TAG) lens were used to generate a rapid 3D laser-scan over a 1 × 1 × 2 μm^3^ volume (Figure 1B, Supplementary Figure 1). Photons are collected on a single-photon-counting avalanche photodiode (APD). The photon arrival time is used to calculate the diffusing particle’s real-time position, which is then used to drive a 3D piezoelectric stage to bring the particle back to the detection center, effectively locking the moving target in focus. 3D-SMART produces a 3D particle trajectory with 1,000 loc s^-1^ with localization precision up to ∼ 20 nm in XY and ∼ 80 nm in Z across the travel range of the piezoelectric stage (∼ 75 × 75 × 50 μm^3^; XYZ).^25^

3D-FASTR was employed to achieve rapid volumetric imaging for environmental contextualization of single virus trajectories. 3D-FASTR uses a two-photon laser-scanning microscope equipped with an electrically tunable lens (ETL). The raster scan pixel dwell time (∼ 1 μs) is significantly shorter than the stage response time (∼ 1 ms), making motion blur negligible. The ETL performs remote focusing of the imaging laser to enable 3D imaging, generating a continuous focal scan across an 8-μm range. By optimizing the scan frequencies, each iteration of the Z focal scan tessellates and tiles to scan the entire volume (Figure 1C, Supplementary Figure 1). At short acquisition times, unsampled voxels will be reconstructed from sampled neighboring voxels, thereby increasing the volumetric imaging rate by 2-4 fold over conventional image-stacking methods.^28^ 3D-TrIm combines 3D-SMART and 3D-FASTR into an integrated platform, where piezoelectric stage readout is registered with the imaging voxel position to overlay highly sampled 3D viral trajectories with volumetric cellular images.

The viral infection process spans a range of temporal scales. While millisecond sampling reveals molecular-level interactions, it loses its utility if observation cannot be extended to infection-relevant timescales. These two timescales are usually at odds, as short timescale observations require strong illumination, which increases photobleaching and limits trajectory lengths. Therefore, it is required to extend tracking duration without sacrificing spatiotemporal precision. Recent advances in fluorescent proteins significantly expanded the photon budget for fluorescent microscopies, including SVT.^29^ StayGold,^30^ a highly photostable green fluorescent protein, has been previously employed in high-temporal-resolution 3D-SVT to extend tracking duration to over an hour.^29^ A previously reported and well-characterized internal virus labeling approach was employed, whereby StayGold is fused to the HIV-1 viral protein R (Vpr) and packaged into budding lentivirus particles (Figure 1D).^31^ Generated particles are pseudotyped with the SARS-CoV-2 D614G spike protein on their surface (CoV2/SG-VLP).^32^ Successful incorporation of Vpr-StayGold and spike protein into single virions was confirmed by tracking CoV2/SG-VLP in the absence of cells (Supplementary Figure 2) and by immunofluorescence experiments (Supplementary Figure 3). The expression level of all cell lines used in this work was assessed through flow cytometry analysis (Supplementary Figure 4a), and the infectivity of generated VLPs across different cell lines was tested by a reporter assay (Supplementary Figure 4b). Similar to previously reported results,^29^ CoV2/SG-VLP significantly extended the 3D-TrIm tracking duration from minutes to over an hour (Supplementary Figure 5).

To preserve fluorescence signal from the fluorescently labeled cells throughout the enhanced tracking duration, a shutter on-and-off feature was incorporated into 3D-TrIm to gate the two-photon imaging laser. The shutter is only turned on for 16 seconds every minute to construct volumetric images, and the volume acquired can be used to construct sequences over the course of the trajectory (1 volume per minute) or accumulated over the duration of an entire trajectory to form a single global volume.

**Figure 1.**
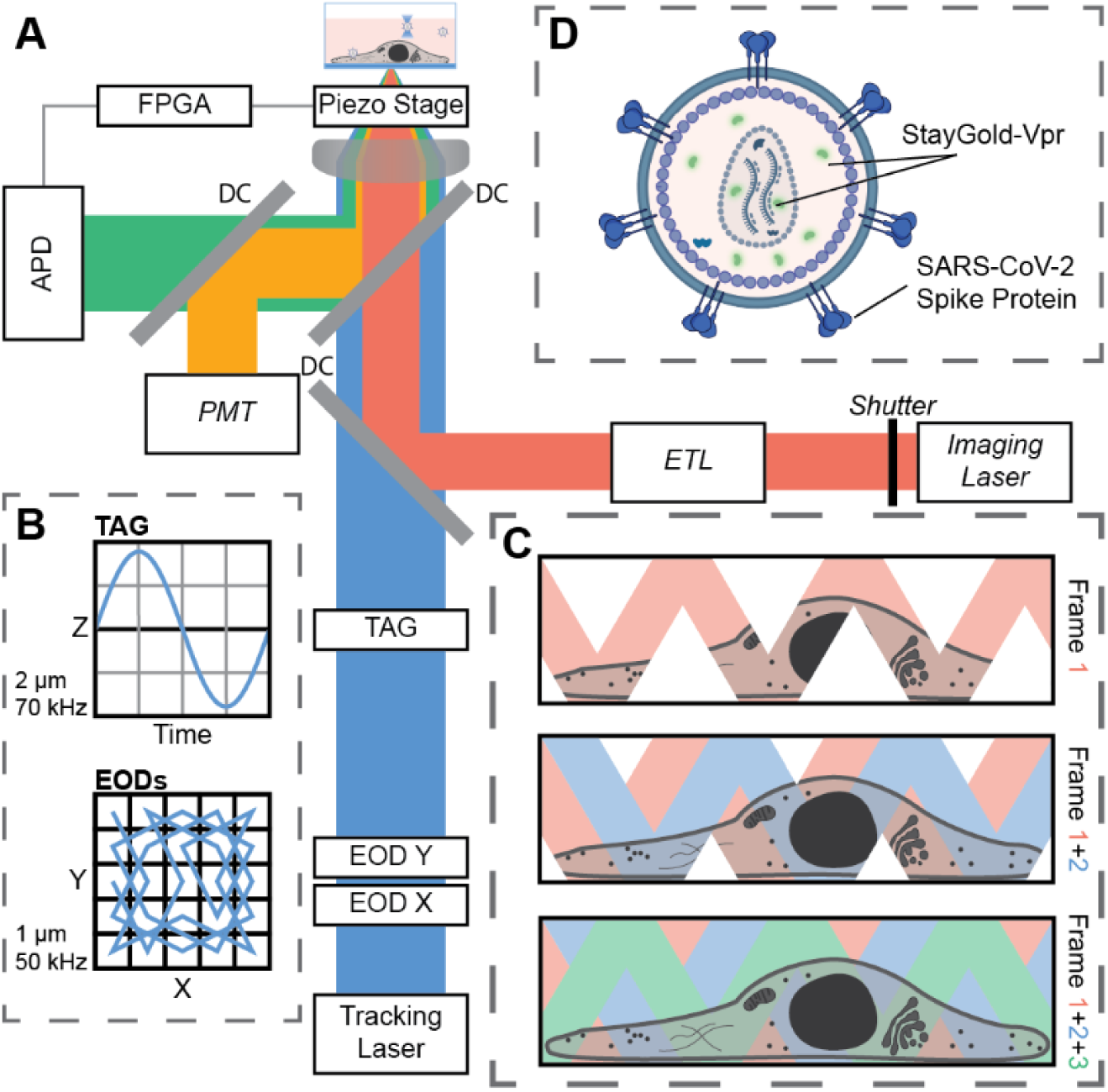
3D-TrIm setup for long-term 3D tracking and imaging of VLPs labelled with StayGold. (**A**) 3D-TrIm setup. Fluorescently labeled VLPs are added to live cells on a coverslip. The sample is placed on a heated sample holder mounted on a piezoelectric stage. (**B**) Overview of 3D-SMART tracking scanning patterns. The EODs generate a fast knight’s tour scanning pattern in a 1 μm × 1 μm range at a frequency of 50 kHz per spot. The TAG lens oscillating at 70 kHz extends the 2D scan axially by ± 1 μm. (**C**) Concept of 3D-FASTR volumetric imaging. An ETL is incorporated into a traditional two-photon LSM to generate a repeatable and tessellated 3D sampling pattern. The entire volume is sampled after a set number of frame times. (**D**) VLPs carrying SARS-CoV-2 spike proteins were internally labelled by encapsulating StayGold-Vpr.

### Viral trajectories with resolvable state transitions captured with extended tracking duration

293T cells naturally express a minimal level of SARS-CoV-2 receptors, such as ACE2 and TMPRSS2, but they can be readily engineered to overexpress those receptors through stable transfection.^33^ To prevent perturbation of actin activity and to offer long-term imaging of the cell morphology, nucleic acid staining (SYTO 61) was chosen instead of receptor-staining or lipophilic staining methods. SVT was performed using 3D-TrIm microscopy and CoV2/SG-VLPs on 293T cells expressing ACE2, the receptor for SARS-CoV-2. Due to the strong photostability of the StayGold labeling, long-term single virus trajectories were captured, covering the early events in SARS-CoV-2 infection. A ∼ 15-minute example trajectory is shown in Figure 2A, which starts with extracellular diffusion for ∼ 20 sec. After multiple attempts and close contact with the cell body, the virus landed and diffused along a cellular protrusion. Although not stained, the high-resolution 3D trace depicts a ∼ 3 μm wavy protrusion ∼ 1 μm away from the cell body (Figure 2A-C), which is similar to previously reported “virus surfing”.^4, 25, 34^ Due to its horizontal orientation and relatively poorer axial localization, this trajectory is not shown as an obvious hollow “ring”. However, from the 3D point cloud color-coded by the density of neighboring points, an uneven distribution of localization still indicates a ring structure (Figure 2B). Further validation was done by simulating a 3D random walk on a cylindrical surface (Supplementary Figure 6, see also Supporting Information for details). A best-fit cylinder gives a radius of 153 ± 60 nm, which is consistent with the observed size of filopodial protrusions. Following ∼170 sec of diffusion on the protrusion, the virus rapidly diffused onto the cell body and underwent directed motion along the cellular membrane. The diffusion coefficient decreased by more than an order of magnitude, from 0.058 ± 0.015 μm^2^/sec to 0.007 ± 0.001 μm^2^/sec, while retaining high directionality near the cellular membrane.

Three quantification methods were applied to analyze the motion of single VLPs at or near the cell surface. These are the virus-to-cell distance, sliding-window-based mean-squared displacement (MSD) analysis, and drift-velocity calculation (see also the Supporting Information for details). An example of the windowed-MSD and virus-to-cell distance calculations is shown in Figure 2C. During the free diffusion stage, the VLP exhibited rapid Brownian motion with a net zero drift velocity (Figure 2C). The distance between the virus and the cell fluctuated, with values near zero indicating close contact with the cell membrane. After binding at ∼20 sec, the diffusion coefficient dropped by roughly an order of magnitude, tracing out a cylindrical structure as described above. Interestingly, despite the cylindrical structure, the average drift velocity remained around zero, suggesting that no favored directionality or directed motion was involved. After the VLP reached the cell membrane (indicated by a virus-to-cell distance near zero), the diffusion coefficient dropped by another order of magnitude. However, the motion appeared to be directed with an overall drift velocity of ∼ 1.2 nm/sec. Compared with the slow diffusion and directed motion on the larger cell membrane, the relatively rapid diffusion of the VLP on the cylindrical protrusion may indicate that different numbers of receptors were bound to the particle.

**Figure 2.**
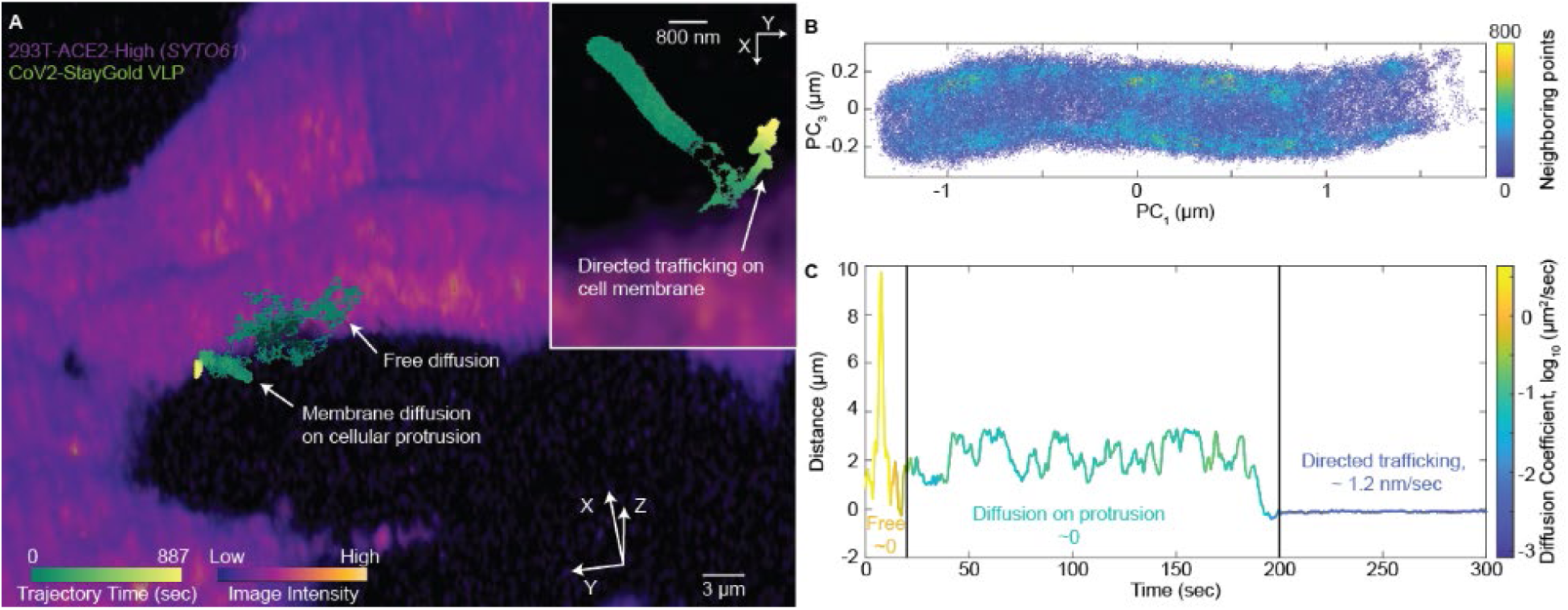
An SVT trajectory with resolvable state transitions. (A) A CoV2/SG-VLP trajectory of viral approach and landing with resolvable state transitions, including extracellular diffusion, landing, diffusion on a cylindrical protrusion, and directed trafficking on the cell membrane (see also Supplementary Video 1). (B) 3D point cloud color-coded by density shows a ring-like structure as previously reported for membrane protusions. (C) The virus-to-cell distance map, color-coded by diffusion coefficient, shows three distinct states. The overall drift velocities are indicated for each state. Virus-to-cell distance trace is a 1-second running mean. Only the first 300 seconds are shown here for ease of visualization.

### Directed viral trafficking observed on the plasma membrane

As previously reported across a broad range of studies, viral surfing along filopodia often results in rapid transport of viral particles to so-called “endocytic hot spots,” after which internalization or fusion occurs.^4, 13–15^ The work presented here demonstrates that viruses undergo further trafficking upon arrival in the cell body through viral surfing. Based on the virus-to-cell distance measurement, this linear trafficking event happens proximal to the cell membrane. This could indicate that directed viral trafficking is a general feature of infection that occurs prior to viral internalization.

Linear trafficking of CoV2/SG-VLPs at the cell membrane was repeatedly observed on 293T cell lines expressing ACE2 receptors. An example 30-minute trajectory of a single CoV2/SG-VLP in 293T-ACE2-TMPRSS2 is shown in Figure 3A. The observed viral membrane trafficking, similar to other well-studied linear trafficking behaviors, is highly directional and shows a “stop-and-go” pattern. The directed viral trafficking tightly matches the contour of the cellular volume (Figure 3B). The trafficking events show a stop-and-go pattern, with drift velocity ranging between 20 and 120 nm/sec (Figure 3E-F, Table 1, and Supplementary Figure 7; see the following section and Methods for detailed information on drift velocity calculation), aligned with previously reported results.^4, 16^ Observed viral trafficking events ranged from tens of seconds to over two minutes (Supplementary Figure 7). Throughout the process, VLP redirection and turning are also observed, which is similar to the pattern of vesicles being trafficked along microtubules or actin filaments, due to the dynamic association and dissociation of motor proteins and the physical constraints of the underlying network structure.^2, 35^ The membrane-proximal viral trafficking reported here and the more widely understood internalized trafficking are readily distinguished by their localization within the cellular volume (Supplementary Figure 8). A distinct difference can be seen in the virus-to-cell distance of membrane-proximal and internalized trafficking (Supplementary Figure 8c). The measured virus-to-cell distances are negative for internalized trafficking throughout the entire trajectory, while distances for membrane trafficking fluctuate around zero for the duration of the trajectory.

For the example described above, after ∼ 28 minutes of membrane-localized directed viral trafficking, the virus-to-cell distance rapidly drops below zero, indicating viral internalization (Figure 3C-E). Notably, the particle exhibits a prolonged period of confined motion at a location spatially separated from the entry point, followed by a second phase of directed motion leading to internalization (Figure 3E-F and Supplementary Figure 7a). During internalization, the viral particle maintains directed motion, suggesting a continuous transport process rather than a discrete transition from membrane trafficking to entry.

Together, these observations indicate that viral entry is not a single-step process, but instead involves multiple stages of surface exploration, including a prolonged period of confined motion followed by directed trafficking toward the final entry site and into the cell. These data suggest that membrane trafficking events could facilitate rapid viral trafficking over large length scales towards endocytic hotspots, where key regulatory proteins and factors are concentrated.

**Figure 3.**
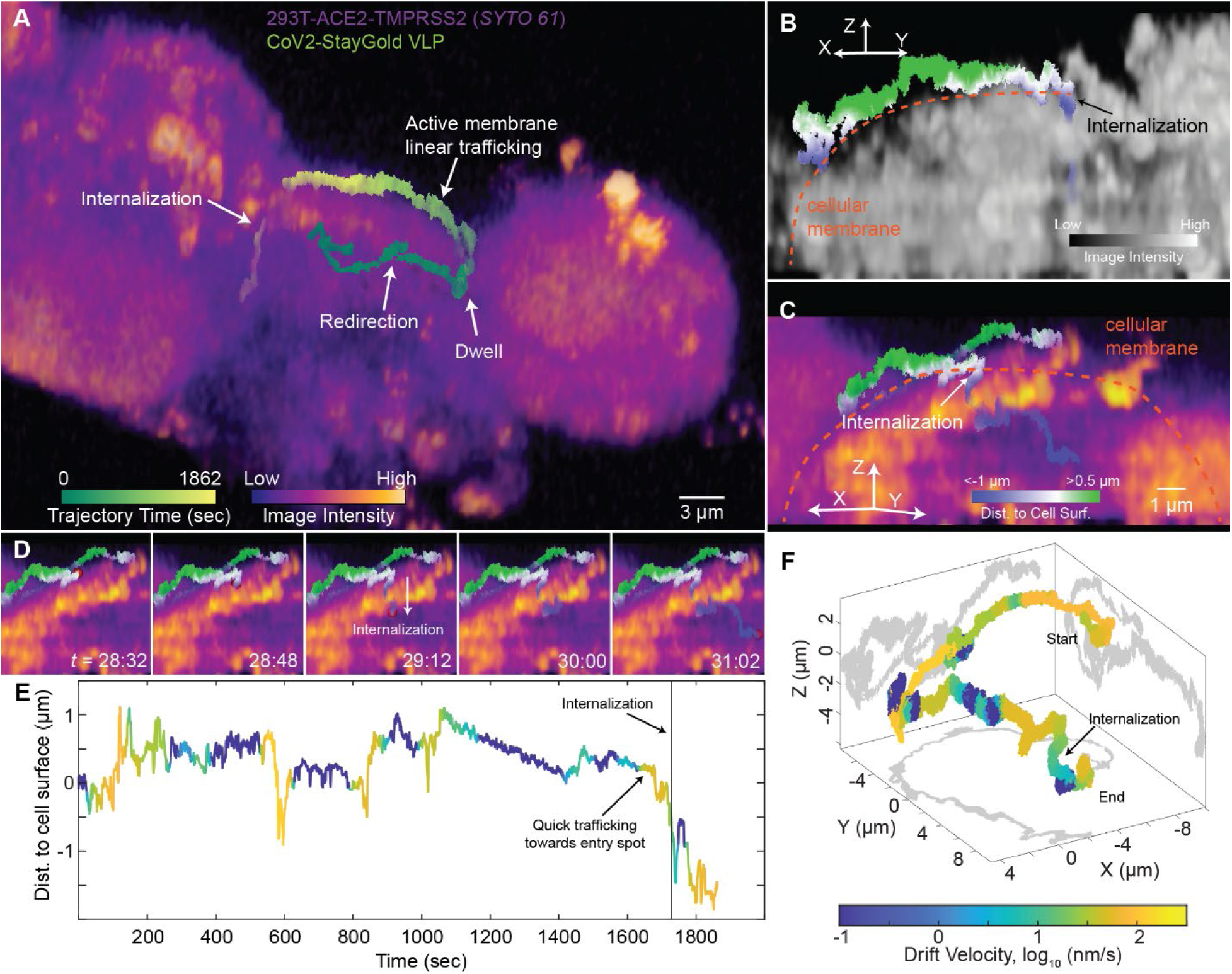
CoV2/SG-VLP actively trafficked on the cell membrane towards an internalization spot. (A) A >30 min trajectory of a CoV2/SG-VLP in 293T-ACE2-TMPRSS2 cells stained with SYTO 61. (B) Active VLP trafficking along the cellular contour. SYTO61 intensity is shown in grayscale. (C) Internalization of the VLP. (D) Five snapshots during membrane trafficking followed by internalization and internalized trafficking. The trajectories in (B-D) are color-coded by the virus-to-cell distance (see also Supplementary Video 2). (E) The virus-to-cell distance map, color-coded by drift velocity, indicating directed trafficking toward the entry site and during internalization. (F) 3D trajectory in (A-D) color-coded by drift velocity.

### Mechanism of viral membrane trafficking

The mechanism of this viral membrane trafficking pattern is investigated from different perspectives, including the presence of ACE2 and TMPRSS2, ACE2 expression level, actin, and tubulin activity. It was first confirmed that viral membrane trafficking is receptor-dependent by repeating the experiment with native 293T cells, which lack ACE2 expression. Under this condition, the majority of trajectories exhibited free diffusion in the extracellular space, with only a few becoming stuck or immobilized near the cell membrane or on the coverslip. No trajectories displaying linear trafficking patterns were observed (data not shown). These results indicate that the observed trafficking events occur after receptor binding, likely involving the transport of virus–receptor complexes along the membrane.

A wide distribution of drift velocities was observed, even within a single VLP trajectory. This can be seen in Figure 4A-C for a CoV2/SG-VLP trajectory on ACE2-expressing 293T cells, where the particle traffics along the membrane and internalizes (Figure 4A-C and F). To quantify linear trafficking events, principal component analysis (PCA)-enhanced MSD analysis was performed in 100-second segments. PCA was performed to identify the direction of maximum variance, corresponding to the direction of trafficking. MSD analysis was then performed along this direction. The minimum trustworthy drift velocity calculated by this algorithm is 0.4 nm/s, which corresponds to a 40 nm displacement over 100 seconds and is determined by the 3D-TrIm tracking localization precision. In this trajectory, a CoV2/SG-VLP was first immobilized on the plasma membrane, then underwent fast trafficking with a ∼ 90 nm/s drift velocity for over 40 seconds. Reaching the end of the fast-trafficking, the VLP was internalized. Compared to the fast membrane trafficking in this specific trajectory, the drift velocity of the intracellular part was slower, around 10 to 20 nm/sec (Figure 4F).

Membrane trafficking events exhibited high directionality on the cellular plasma membrane, supporting the hypothesis that they are actin-aided. Previous research suggested the existence of viral motility that is actin-dependent and microtubule-independent.^36^ Other works have also observed colocalization of SARS-CoV-2 viral particles with actin.^37^ To test the hypothesis that membrane-localized directed viral trafficking is actin dependent, ACE2-expressing 293T cells were pretreated with SMIFH2 prior to tracking experiments, which is a small molecule inhibitor of formin FH2 domains, inhibiting the polymerization-depolymerization cycle of actin filaments while also perturbing myosin activity.^38–40^ After SMIFH2 treatment on 293T cells with a high level of ACE2 expression (293T-ACE2-high. See Supplementary Figure 4), a significant drop in drift velocity is observed (Figure 4D, Table 1). An example of an inhibited trafficking trajectory is shown in Figure 4E, where the drift velocities were reduced to ∼ 1-10 nm/sec. Similar results were also observed with CK-666 treatment (Figure 4D, Table 1), an Arp2/3 complex inhibitor that inhibits actin polymerization.^41^ Notably, the reduction in drift velocity was less pronounced in the CK-666-treated group than in the SMIFH2-treated group. This difference may reflect the broader inhibitory effects of SMIFH2, which has been reported to perturb not only formin-mediated actin assembly but also myosin activity, whereas CK-666 more specifically targets Arp2/3-dependent actin polymerization. These results suggest that the trafficking events are strongly correlated with actin dynamics, while also suggesting that multiple components of the actin machinery may contribute to this process, including actin polymerization–depolymerization and myosin activity.

The role of tubulin in these membrane trafficking events was also tested by introducing nocodazole to perturb microtubule polymerization.^42^ After nocodazole treatment using a previously described protocol,^43^ linear membrane trafficking events were still observed, and drift velocity was only minorly impacted compared to the untreated group. This affirmed that membrane trafficking is microtubule-independent and further excluded the possibility that the observed membrane-proximal trafficking is simply near-membrane endosomal trafficking. It was also observed that the velocity of membrane trafficking is dependent on the expression level of ACE2. The different expression levels of two 293T-ACE2 cell lines were verified through both flow cytometry and infection assay (Supplementary Figure 4). The drift velocity from 293T cells with a low ACE2 expression level (293T-ACE2-low) is significantly slower compared to 293T-ACE2 cells with higher ACE2 receptor expression level (Figure 4D, Table 1). Interestingly, CoV2/SG-VLP membrane trafficking events were also observed on cells coexpressing ACE2 and TMPRSS2, but with slightly slower drift velocity. 293T-ACE2-TMPRSS2 cells were also found to be less sensitive to SMIFH2 treatment (Supplementary Figure 9a). One possible explanation is that different infection pathways are engaged, which may not strongly depend on actin activity.

**Figure 4.**
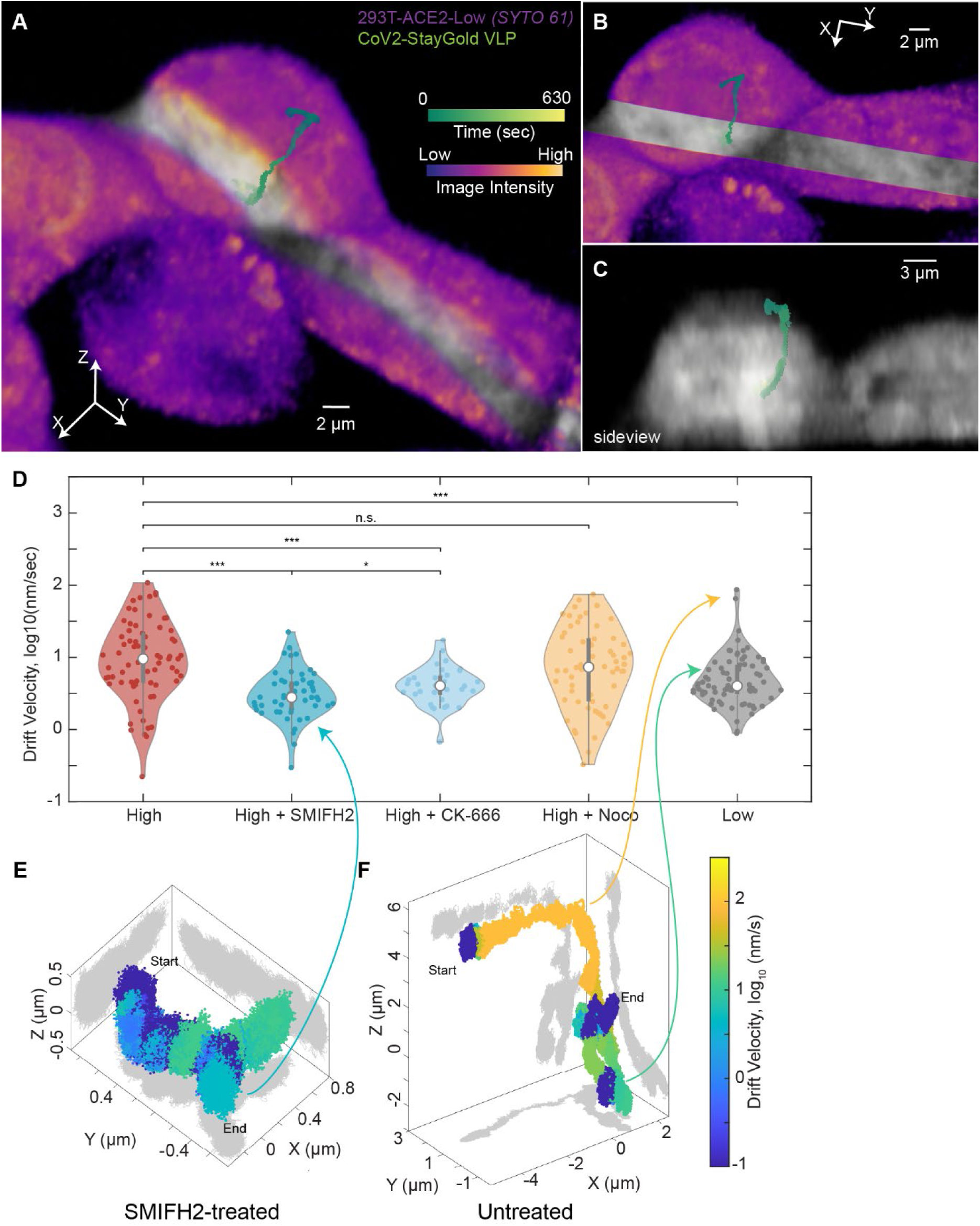
Membrane viral trafficking is regulated by actin activity and expression level of receptors. (A) A 10-minute CoV2/SG-VLP trajectory on 293T-ACE2 cells stained with SYTO 61. Two slice planes were used to section the YZ plane and create a channel that renders the interior volume section semitransparent. The VLP undergoes membrane trafficking, followed by internalization. (B) Top-down view. (C) Side view showing internalization. (D) Violin plot of the drift velocity, expressed as log_10_ (nm/sec), calculated from 100-sec segments. *N* = 70, 51, 34, 60, 76, respectively. n.s.: no significance, *: *p*<0.05, **: *p*<0.01, ***: *p*<0.001, Kolmogorov-Smirnov test. (E) 3D SVT trajectory of a CoV2/SG-VLP on SMIFH2 treated 293T-ACE2 cells, color-coded by drift velocity. (F) 3D trajectory in (A-C) color-coded by drift velocity. Three arrows in (E-F) point to the value of three representative drift velocities in the violin plot.

**Table 1:**
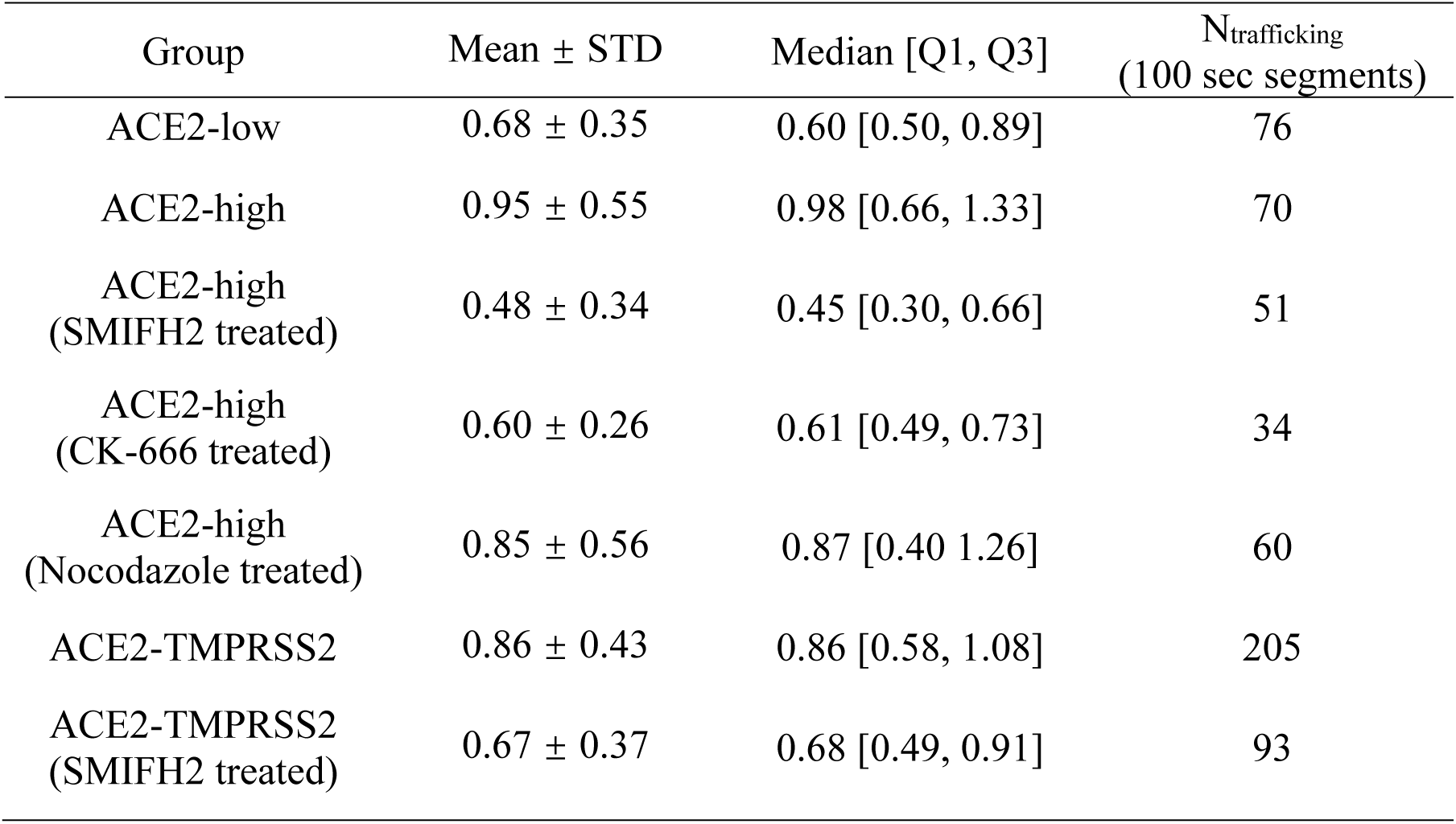
Distribution of drift velocity of StayGold/SG-VLP in different cells. All values are expressed as log_10_-transformed drift velocity (nm/sec).

## Discussion

We would like to emphasize the technical challenges in capturing and characterizing this type of membrane-trafficking motility. To resolve such motion, an SVT method must provide sufficient spatiotemporal resolution in 3D while maintaining a sufficiently long observation window. The inherently volumetric capability of the 3D-TrIm microscope provided the necessary instrumental advantage, particularly in capturing viral dynamics along a large axial scale during these membrane trafficking events. Meanwhile, the use of StayGold labelled VLPs substantially extended the achievable tracking duration without compromising tracking performance, further facilitating the successful observation of this long-lasting motility. Moreover, low photon dose of the tracking microscope also minimized the amount of photons impinging upon the live cells, greatly reducing photobleaching compared to other volumetric imaging methods, such as spinning-disk confocal microscopy.

Based on the experimental results presented above via long-term and high-resolution 3D-SVT using 3D-TrIm, membrane trafficking shows a high degree of similarity to previously described “viral surfing” on membrane protrusions.^4^ Both motilities require cognate Env-receptor interaction and are regulated by actin activity. Instead of “surfing” along actin-rich protrusions, membrane trafficking can happen on membrane-peripheral actin-rich structures, such as the actin cortex. Based on the previously proposed mechanism of protrusion-based viral surfing, membrane trafficking may be critical for relocating the virus towards the endocytic hotspot. This process involves virus attachment, receptor recruitment, the establishment of a link to the underlying actin cytoskeleton, and actin-aided transport. The directed motion could be similarly powered by retrograde F-actin flow with myosin II or through the polymerization and depolymerization of the actin cytoskeleton.

The mechanism of how viruses initiate viral surfing and membrane trafficking remains largely uncharacterized.^3^ Zhang et al visualized that SARS-CoV-2 can regulate and utilize filopodia in two patterns, “surfing” and “grabbing”,^16^ which suggested its ability to hijack actin. Previous research has suggested that ACE2 receptors are associated with actin through multiple molecules. Fujimoto et al revealed cytoplasmic actin as a strong interactor of the omicron receptor binding domain (RBD).^44^ Ogunlade et al. confirmed the interaction between the actin-bundling protein Fascin-1 and ACE2.^45^ There are also multiple reports that ACE2 binds integrins, which are adhesion molecules, and can regulate their signaling.^46–48^ Also, signaling from Env-ACE2 interactions could regulate and remodel the actin cytoskeleton through multiple pathways. It has been demonstrated that SARS-CoV-2 infection could induce the upregulation of the actin cytoskeleton via Rho-family GTPases such as cell division cycle 42 (Cdc42),^16^ rearrangement of actin under the regulation of the protein kinase N (PKN) and P21-activated kinase 1 (PAK1),^37, 49^ and dephosphorylation of Ezrin/Radixin/Moesin (ERM) proteins to reduce the stiffness of cortical actin.^50^

How the expression level of ACE2 affects viral surfing on the cell membrane remains to be determined. Higher ACE2 expression could lead to stronger signaling due to multivalent binding. Higher ACE2 expression levels could also lead to more effective interaction between the virus-receptor complex and actin, increasing the chance of active trafficking. On the other hand, the higher degree of multivalent binding of SARS-CoV-2 can also possibly result in stronger signaling and higher degree of actin remodeling, thus faster membrane trafficking.

**Figure 5.**
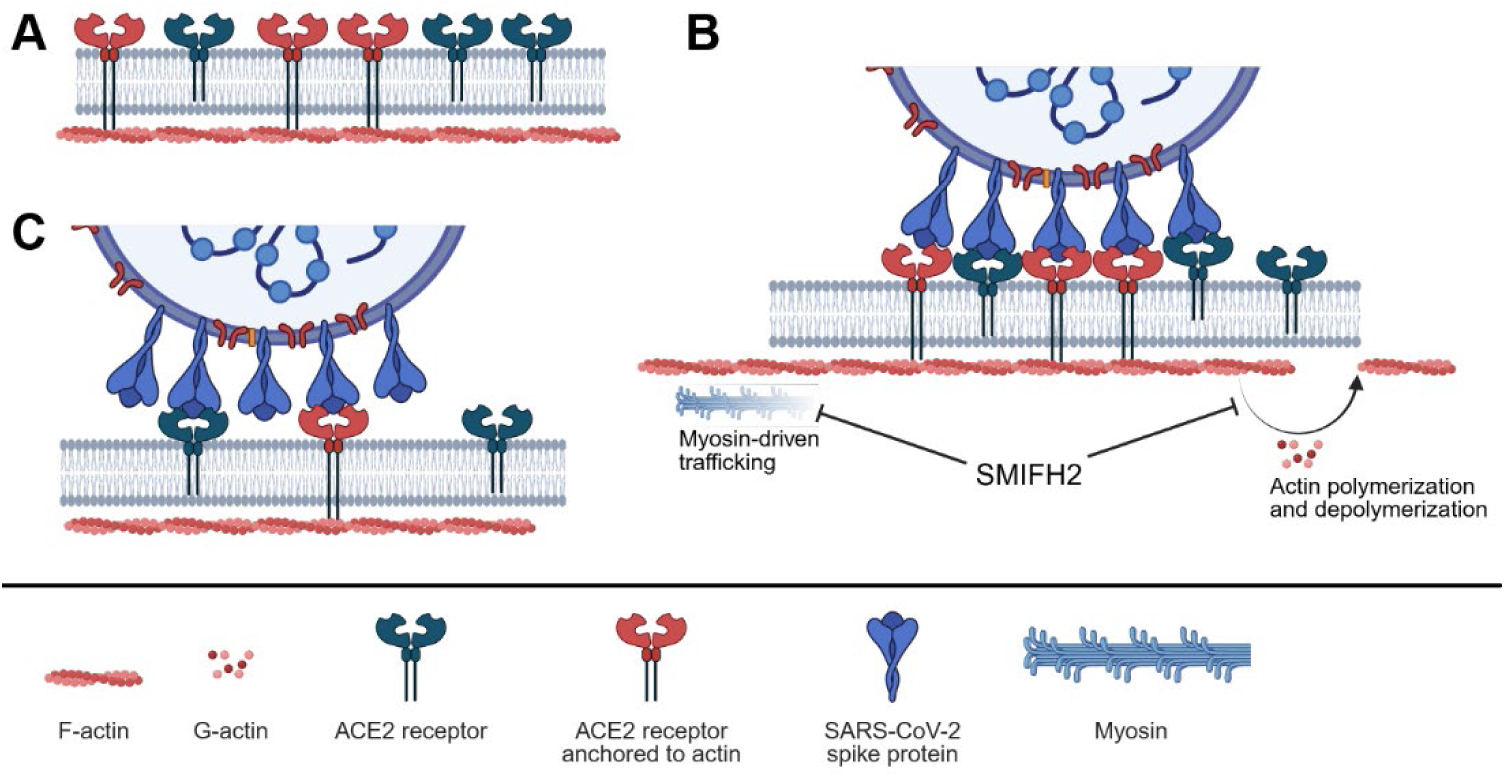
Proposed mechanisms of viral trafficking on the plasma membrane. (A) In cells expressing ACE2 receptors, two populations of receptors exist: one actively interacts with actin, and another does not. (B) Upon viral binding, multivalent interactions occur between spike proteins and receptors. The binding of actin-interacting ACE2 could increase the affinity between the virus and actin or downstream signalling, resulting in more active trafficking events. Trafficking events are driven by either myosin-based transport or actin polymerization and depolymerization, and these processes can be inhibited by SMIFH2 or CK-666 treatment. (C) In cells with lower ACE2 expression, fewer spike-receptor interactions occur, reducing the affinity/signalling and slowing down trafficking.

Viral surfing along actin-rich protrusions has been widely observed across diverse virus–cell systems and is considered a general mechanism for facilitating viral entry.^4, 13–15^ Building on this established paradigm, our observations suggest that actin-assisted trafficking may not be limited to protrusive structures. Instead, viruses may similarly exploit membrane-associated actin networks to undergo directed motion along the plasma membrane. While demonstrated here for SARS-CoV-2, this membrane trafficking behavior is likely to represent a more general strategy across virus–cell models.

This work also highlights the power of high-speed 3D SVT as a tool to resolve viral entry with high spatiotemporal resolution in three dimensions. For example, in both Figure 3 and Figure 4, viral particles continue to exhibit a linear trafficking mode during internalization into the cells. However, it remains challenging to draw statistically robust conclusions for rare and transient events, such as viral entry, due to the limited throughput of active-feedback based approaches. Future work could address this limitation by incorporating event-driven algorithms into the current workflow.^51^ For example, image-based screening methods could be used to identify particles exhibiting targeted behaviors, followed by real-time switching to active-feedback tracking for high-speed SVT. Such hybrid approaches may enable efficient detection and detailed characterization of rare infection events.

In addition, integrating functional probes into this platform could further elucidate the mechanisms underlying viral internalization. For instance, pH-sensitive reporters could be used to monitor endosomal acidification,^52^ while membrane labeling strategies could provide direct visualization of membrane remodeling during entry.^53^ Coupled with long-term 3D tracking, such approaches may enable simultaneous correlation of viral motion with biochemical and structural transitions, providing a more complete picture of the viral entry process.

## Conclusion

Taken together, given the ubiquitous distribution of actin, it is not surprising that viruses can hijack and regulate actin during viral infections. Actin-rich structures, especially protrusions such as filopodia, have often been used by viruses to rapidly approach the cell body. Here, we reveal the novel process in which viruses induce rapid and directed trafficking on the plasma membrane to explore the cell and move to the optimal entry spot. This membrane trafficking motility was found to be a post-binding event, and its magnitude is positively correlated with the receptor expression level and the actin activity. These data suggest that viruses can hijack and regulate actin in host cells and could ride the cell membrane, potentially through the actin cortex, to rapidly explore the cell and relocate to an optimal entry site.

## Materials and Methods

### 3D-TrIm Overview

The 3D-TrIm instrument design is shown in Supplementary Figure 1 and consists of 3D-SMART tracking excitation optics and 3D-FASTR imaging excitation optics coupled through a commercial confocal microscope (Zeiss LSM 410, modified by LSM Tech), piezoelectric stage and microscope objective to join both setups together. The microscope is controlled by custom LabVIEW code.

### Sample Preparation and Experimental Setup

At 24 h prior to live-cell single-virus tracking experiments, cells were plated in complete DMEM at 8 × 10^5^ cells per well in a 6-well plate with an autoclaved glass coverslip (VWR, no. CLS-1760-025) to ensure 80% confluency on the day of the experiment. The coverslips were treated with poly-lysine for better adhesion of 293T cells and to avoid stacking of cells in multiple layers. Cells were stained with nucleic acid stain (SYTO 61, ThermoFisher, no. S11343) by incubating with 500 nM SYTO 61 in complete DMEM for 30 min at 37 °C and 5% CO_2_ followed by careful wash with 1×PBS. After staining, the coverslip was transferred to HEPES pH 7.4 buffered solution (live cell imaging solution, LCIS, ThermoFisher, no. A14291DJ) in a custom-built sample holder and positioned on a heated stage (37 °C). All experiments were performed using live cells.

Cell lines used in this paper includes 293T/17 (Duke Cell Culture Facility, ATCC# CRL-11268), 293T-hACE2-low (by transducing 293T/17 with lentivirus encoding hACE2), 293T-hACE2-high (BEI Resource, NR-52512), 293T-ACE2.TMPRSS2 (BEI Resource, NR-55293, derived from NR-52512 mentioned above). Live cell volumetric imaging and real-time viral tracking were performed on cells prepared as described above. CoV2-StayGold VLP was added to cells on a heated stage to initiate the reaction to a final concentration of 1.9 × 10^6^ TU ml^−1^, which corresponds to multiplicity of infection ∼1. The average excitation power at the focus was 180 nW and 7.5 mW for the 488-nm (tracking) and 780-nm (imaging) beams, respectively.

For SMIFH2 treatment, SMIFH2 (Sigma-Aldrich, S4826) was prepared into 50 mM stock in DMSO, stored in aliquots at −20°C. Before 3D-TrIm experiments, cells were treated with 25 μM SMFIH2 for 1 hr. During the experiment, LCIS with an additional 12.5 μM SMIFH2 was used to maintain the inhibition.

For CK-666 treatment, CK-666 (Sigma-Aldrich, SML0006) was prepared into 10 mM stock in DMSO, stored in aliquots at −20°C. Before 3D-TrIm experiments, cells were treated with 100 μM CK-666 for 1 hr. During the experiment, LCIS with an additional 100 μM CK-666 was used to maintain the inhibition.

For nocodazole treatment, nocodazole (Sigma-Aldrich, M1404) was prepared into 10 mM stock in DMSO, stored in aliquots at −20°C. Before 3D-TrIm experiments, cells were treated with 30 μM nocodazole for 1 hr. During the experiment, LCIS with an additional 30 μM nocodazole was used to maintain the inhibition.

### Normalization of Anisotropic 3D Localization Precision

Due to the difference in lateral (XY) and axial (Z) precision, the localizations were normalized before processing PCA. Briefly, variances of XYZ coordinates in 200 ms segments were calculated and the mean value is used to represent the variance throughout the entire window. Z axis is normalized based on the ratio between variance in XY and variance in Z.

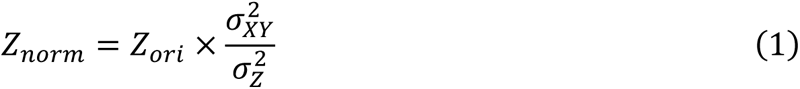

where *Z_ori_* is the original Z axis, *Z_norm_* is the Z axis after normalization, *σ_XY_* is the lateral precision, and *σ_Z_* is the axial precision.

### PCA-Enhanced MSD Analysis for Linear Trafficking

To improve the characterization of particles undergoing directed or linear trafficking, we applied a principal component analysis (PCA)-based approach to enhance mean square displacement (MSD) analysis. Briefly, PCA identifies the directions (principal components, for 3D trajectories, PC1-3) along which the variance in the data is maximized and projects the data onto these directions (score 1-3). Specifically, the eigenvectors (coefficients) obtained from PCA represent the new orthogonal coordinate system, where the first coefficient corresponds to the direction of maximal variance in the trajectory. Explained variance (explained%) can be used to determine the contribution of three PCs, differentiating anisotropic and isotropic motions.

When PC1 explained% is over 80 %, 1D MSD was performed along score 1. When PC1 explained% is less than 80% while PC3 explained% is less than 25% of the variance, 2D MSD was performed on score 1 and 2. When PC3 explained% is over 25%, 3D MSD was performed.

## Acknowledgements

We thank the Duke Cell Culture Facility for access to cell lines used in this study. We gratefully acknowledge the Duke Light Microscopy Core Facility and Brittney Ferrari from the Duke Cancer Institute Flow Cytometry Facility for their support and assistance in this work. FACS was performed using Beckman Coulter Astrios cell sorter in the Duke Cancer Institute Flow Cytometry Facility at Duke University, Durham, NC, which is supported by the NCI Cancer Center Support Grant (CCSG) award number P30CA014236. Some schematics were created with BioRender.com. The following reagents were obtained through BEI Resources, NIAID, NIH: (1) *Homo sapiens* Embryonic Kidney Epithelial Cells Expressing Transmembrane Protease, Serine 2 and Human Angiotensin-Converting Enzyme 2 [293T-ACE2.TMPRSS2 (mCherry)], NR-55293. (2) Human Embryonic Kidney Cells (HEK-293T) Expressing Human Angiotensin-Converting Enzyme 2, HEK-293T-hACE2 Cell Line, NR-52512. (3) Polyclonal Anti-SARS-Related Coronavirus 2 Spike Glycoprotein (IgG, Rabbit), NR-52947.

## Author contributions

Y.L. and K.D.W. conceptualized the study. Y.L., J.E., C.J. and K.D.W. were responsible for the methodology. Y.L., X.L., J.E., and H.L. performed investigations. Y.L., X.L., and K.D.W. curated the data. K.D.W. was responsible for funding acquisition. Y.L. and K.D.W. wrote the original draft, and all authors reviewed and edited the manuscript.

## Data and materials availability

All data needed to evaluate the conclusions in the paper are present in the paper and/or the Supplementary Materials. Additional data related to this paper may be requested from the authors.

## Reference

(1) Dominguez, R.; Holmes, K. C. Actin structure and function. Annual review of biophysics 2011, 40 (1), 169–186.

(2) Chenouard, N.; Xuan, F.; Tsien, R. W. Synaptic vesicle traffic is supported by transient actin filaments and regulated by PKA and NO. Nat. Commun. 2020, 11 (1), 5318.

(3) Taylor, M. P.; Koyuncu, O. O.; Enquist, L. W. Subversion of the actin cytoskeleton during viral infection. Nat. Rev. Microbiol. 2011, 9 (6), 427–439.

(4) Lehmann, M. J.; Sherer, N. M.; Marks, C. B.; Pypaert, M.; Mothes, W. Actin-and myosin-driven movement of viruses along filopodia precedes their entry into cells. The Journal of cell biology 2005, 170 (2), 317–325.

(5) Kloc, M.; Uosef, A.; Wosik, J.; Kubiak, J. Z.; Ghobrial, R. M. Virus interactions with the actin cytoskeleton—what we know and do not know about SARS-CoV-2. Arch. Virol 2022, 167 (3), 737–749.

(6) Walsh, D.; Naghavi, M. H. Exploitation of cytoskeletal networks during early viral infection. Trends in microbiology 2019, 27 (1), 39–50.

(7) Courtemanche, N. Mechanisms of formin-mediated actin assembly and dynamics. Biophysical reviews 2018, 10 (6), 1553–1569.

(8) Molinie, N.; Gautreau, A. The Arp2/3 regulatory system and its deregulation in cancer. Physiological reviews 2018, 98 (1), 215–238.

(9) Svitkina, T. M. Actin cell cortex: structure and molecular organization. Trends in cell biology 2020, 30 (7), 556–565.

(10) Sauvanet, C.; Wayt, J.; Pelaseyed, T.; Bretscher, A. Structure, regulation, and functional diversity of microvilli on the apical domain of epithelial cells. Annu. Rev. Cell Dev. Biol. 2015, 31 (1), 593–621.

(11) Tojkander, S.; Gateva, G.; Lappalainen, P. Actin stress fibers–assembly, dynamics and biological roles. J. Cell Sci. 2012, 125 (8), 1855–1864.

(12) Mattila, P. K.; Lappalainen, P. Filopodia: molecular architecture and cellular functions. Nature reviews Molecular cell biology 2008, 9 (6), 446–454.

(13) Burckhardt, C. J.; Greber, U. F. Virus movements on the plasma membrane support infection and transmission between cells. PLoS pathogens 2009, 5 (11), e1000621.

(14) Schelhaas, M.; Ewers, H.; Rajamäki, M.-L.; Day, P. M.; Schiller, J. T.; Helenius, A. Human papillomavirus type 16 entry: retrograde cell surface transport along actin-rich protrusions. PLoS pathogens 2008, 4 (9), e1000148.

(15) Heusermann, W.; Hean, J.; Trojer, D.; Steib, E.; von Bueren, S.; Graff-Meyer, A.; Genoud, C.; Martin, K.; Pizzato, N.; Voshol, J.; et al. Exosomes surf on filopodia to enter cells at endocytic hot spots, traffic within endosomes, and are targeted to the ER. J. Cell Biol. 2016, 213 (2), 173–184. DOI: 10.1083/jcb.201506084.

(16) Zhang, Y.; Zhang, X.; Li, Z.; Zhao, W.; Yang, H.; Zhao, S.; Tang, D.; Zhang, Q.; Li, Z.; Liu, H. Single particle tracking reveals SARS-CoV-2 regulating and utilizing dynamic filopodia for viral invasion. Science Bulletin 2023, 68 (19), 2210–2224.

(17) Seyran, M.; Takayama, K.; Uversky, V. N.; Lundstrom, K.; Palù, G.; Sherchan, S. P.; Attrish, D.; Rezaei, N.; Aljabali, A. A.; Ghosh, S. The structural basis of accelerated host cell entry by SARS-CoV-2. The FEBS journal 2021, 288 (17), 5010–5020.

(18) Chang, K.; Baginski, J.; Hassan, S. F.; Volin, M.; Shukla, D.; Tiwari, V. Filopodia and viruses: an analysis of membrane processes in entry mechanisms. Frontiers in microbiology 2016, 7, 300.

(19) Du, L.; Ma, A.-X.; Chen, J.-X.; Li, J.; Wang, Z.-G.; Pang, D.-W. Revealing the Role of Actin Cytoskeleton in Herpes Simplex Virus Type 1–Endosome Membrane Fusion by Single-Virus Tracking. ACS nano 2025.

(20) Liu, S.-L.; Wang, Z.-G.; Xie, H.-Y.; Liu, A.-A.; Lamb, D. C.; Pang, D.-W. Single-Virus Tracking: From Imaging Methodologies to Virological Applications. Chem. Rev. 2020, 120 (3), 1936–1979. DOI: 10.1021/acs.chemrev.9b00692.

(21) Christie, S. M.; Tada, T.; Yin, Y.; Bhardwaj, A.; Landau, N. R.; Rothenberg, E. Single-virus tracking reveals variant SARS-CoV-2 spike proteins induce ACE2-independent membrane interactions. Sci. Adv. 2022, 8 (49), eabo3977.

(22) Shen, H.; Tauzin, L. J.; Baiyasi, R.; Wang, W.; Moringo, N.; Shuang, B.; Landes, C. F. Single Particle Tracking: From Theory to Biophysical Applications. Chem. Rev. 2017, 117 (11), 7331–7376. DOI: 10.1021/acs.chemrev.6b00815.

(23) Hou, S.; Johnson, C.; Welsher, K. Real-Time 3D Single Particle Tracking: Towards Active Feedback Single Molecule Spectroscopy in Live Cells. Molecules 2019, 24 (15). DOI: 10.3390/molecules24152826.

(24) Van Heerden, B.; Vickers, N. A.; Krüger, T. P.; Andersson, S. B. Real-Time Feedback-Driven Single-Particle Tracking: A Survey and Perspective. Small 2022, 18 (29), 2107024.

(25) Johnson, C.; Exell, J.; Lin, Y.; Aguilar, J.; Welsher, K. D. Capturing the start point of the virus–cell interaction with high-speed 3D single-virus tracking. Nat. Methods 2022, 19 (12), 1642–1652. DOI: 10.1038/s41592-022-01672-3.

(26) Hou, S.; Exell, J.; Welsher, K. Real-time 3D single molecule tracking. Nat. Commun. 2020, 11 (1), 3607. DOI: 10.1038/s41467-020-17444-6.

(27) Lin, Y.; Lin, H.; Welsher, K. D. Super-Resolving Particle Diffusion Heterogeneity in Porous Hydrogels via High-Speed 3D Active-Feedback Single-Particle Tracking Microscopy. Small 2025, 21 (40), e05319.

(28) Johnson, C.; Exell, J.; Kuo, J.; Welsher, K. Continuous focal translation enhances rate of point-scan volumetric microscopy. Opt. Express 2019, 27 (25), 36241–36258.

(29) Lin, Y.; Exell, J.; Lin, H.; Zhang, C.; Welsher, K. D. Hour-long, kilohertz sampling Rate three-dimensional single-virus tracking in live cells enabled by StayGold fluorescent protein fusions. J. Phys. Chem. B 2024.

(30) Hirano, M.; Ando, R.; Shimozono, S.; Sugiyama, M.; Takeda, N.; Kurokawa, H.; Deguchi, R.; Endo, K.; Haga, K.; Takai-Todaka, R. A highly photostable and bright green fluorescent protein. Nat. Biotechnol. 2022, 1–11.

(31) Desai, T. M.; Marin, M.; Sood, C.; Shi, J.; Nawaz, F.; Aiken, C.; Melikyan, G. B. Fluorescent protein-tagged Vpr dissociates from HIV-1 core after viral fusion and rapidly enters the cell nucleus. Retrovirology 2015, 12 (1), 1–20.

(32) Plante, J. A.; Liu, Y.; Liu, J.; Xia, H.; Johnson, B. A.; Lokugamage, K. G.; Zhang, X.; Muruato, A. E.; Zou, J.; Fontes-Garfias, C. R. Spike mutation D614G alters SARS-CoV-2 fitness. Nature 2021, 592 (7852), 116–121.

(33) Hoffmann, M.; Kleine-Weber, H.; Schroeder, S.; Krüger, N.; Herrler, T.; Erichsen, S.; Schiergens, T. S.; Herrler, G.; Wu, N.-H.; Nitsche, A. SARS-CoV-2 cell entry depends on ACE2 and TMPRSS2 and is blocked by a clinically proven protease inhibitor. cell 2020. 181 (2), 271–280. e278.

(34) Welsher, K.; Yang, H. Multi-resolution 3D visualization of the early stages of cellular uptake of peptide-coated nanoparticles. Nature nanotechnology 2014, 9 (3), 198–203.

(35) D’Souza, A. I.; Grover, R.; Monzon, G. A.; Santen, L.; Diez, S. Vesicles driven by dynein and kinesin exhibit directional reversals without regulators. Nat. Commun. 2023, 14 (1), 7532.

(36) Vaughan, J. C.; Brandenburg, B.; Hogle, J. M.; Zhuang, X. Rapid actin-dependent viral motility in live cells. Biophys. J. 2009, 97 (6), 1647–1656.

(37) Swain, J.; Merida, P.; Rubio, K.; Bracquemond, D.; Neyret, A.; Aguilar-Ordoñez, I.; Günther, S.; Barreto, G.; Muriaux, D. F-actin nanostructures rearrangements and regulation are essential for SARS-CoV-2 particle production in host pulmonary cells. Iscience 2023, 26 (8).

(38) Rizvi, S. A.; Neidt, E. M.; Cui, J.; Feiger, Z.; Skau, C. T.; Gardel, M. L.; Kozmin, S. A.; Kovar, D. R. Identification and characterization of a small molecule inhibitor of formin-mediated actin assembly. Chemistry & biology 2009, 16 (11), 1158–1168.

(39) Isogai, T.; Van Der Kammen, R.; Innocenti, M. SMIFH2 has effects on Formins and p53 that perturb the cell cytoskeleton. Scientific reports 2015, 5 (1), 9802.

(40) Nishimura, Y.; Shi, S.; Zhang, F.; Liu, R.; Takagi, Y.; Bershadsky, A. D.; Viasnoff, V.; Sellers, J. R. The formin inhibitor SMIFH2 inhibits members of the myosin superfamily. J. Cell Sci. 2021, 134 (8), jcs253708.

(41) Hetrick, B.; Han, M. S.; Helgeson, L. A.; Nolen, B. J. Small molecules CK-666 and CK-869 inhibit actin-related protein 2/3 complex by blocking an activating conformational change. Chemistry & biology 2013, 20 (5), 701–712.

(42) Vasquez, R. J.; Howell, B.; Yvon, A.; Wadsworth, P.; Cassimeris, L. Nanomolar concentrations of nocodazole alter microtubule dynamic instability in vivo and in vitro. Molecular biology of the cell 1997, 8 (6), 973–985.

(43) Yea, C.; Dembowy, J.; Pacione, L.; Brown, M. Microtubule-mediated and microtubule-independent transport of adenovirus type 5 in HEK293 cells. Journal of virology 2007, 81 (13), 6899–6908.

(44) Fujimoto, A.; Kawai, H.; Kawamura, R.; Kitamura, A. Interaction of Receptor-Binding Domain of the SARS-CoV-2 Omicron Variant with hACE2 and Actin. Cells 2024, 13 (16), 1318.

(45) Ogunlade, B.; Guidry, J. J.; Mukerjee, S.; Sriramula, S.; Lazartigues, E.; Filipeanu, C. M. The actin bundling protein Fascin-1 as an ACE2-accessory protein. Cellular and Molecular Neurobiology 2022, 42 (1), 255–263.

(46) Clarke, N. E.; Fisher, M. J.; Porter, K. E.; Lambert, D. W.; Turner, A. J. Angiotensin converting enzyme (ACE) and ACE2 bind integrins and ACE2 regulates integrin signalling. PloS one 2012, 7 (4), e34747.

(47) Lin, Q.; Keller, R. S.; Weaver, B.; Zisman, L. S. Interaction of ACE2 and integrin β1 in failing human heart. Biochimica et Biophysica Acta (BBA)-Molecular Basis of Disease 2004, 1689 (3), 175–178.

(48) Gressett, T. E.; Nader, D.; Robles, J. P.; Buranda, T.; Kerrigan, S. W.; Bix, G. Integrins as therapeutic targets for SARS-CoV-2. Frontiers in Cellular and Infection Microbiology 2022, 12, 892323.

(49) Liu, M.; Lu, B.; Li, Y.; Yuan, S.; Zhuang, Z.; Li, G.; Wang, D.; Ma, L.; Zhu, J.; Zhao, J. P21-activated kinase 1 (PAK1)-mediated cytoskeleton rearrangement promotes SARS-CoV-2 entry and ACE2 autophagic degradation. Signal transduction and targeted therapy 2023, 8 (1), 385.

(50) Do, T. L.; Tachibana, K.; Yamamoto, N.; Ando, K.; Isoda, T.; Kihara, T. Interaction of SARS-CoV-2 Spike protein with ACE2 induces cortical actin modulation, including dephosphorylation of ERM proteins and reduction of cortical stiffness. Human Cell 2024, 38 (1), 3.

(51) Shi, Y.; Tabet, J. S.; Milkie, D. E.; Daugird, T. A.; Yang, C. Q.; Ritter, A. T.; Giovannucci, A.; Legant, W. R. Smart lattice light-sheet microscopy for imaging rare and complex cellular events. Nat. Methods 2024, 21 (2), 301–310.

(52) Fares, P.; Duhaini, M.; Tripathy, S. K.; Srour, A.; Kondapalli, K. C. Acidic pH of early endosomes governs SARS-CoV-2 transport in host cells. J. Biol. Chem. 2025, 301 (2), 108144.

(53) Miyauchi, K.; Kim, Y.; Latinovic, O.; Morozov, V.; Melikyan, G. B. HIV enters cells via endocytosis and dynamin-dependent fusion with endosomes. Cell 2009, 137 (3), 433–444.

