## Supplementary Information for "High-speed 3D single-virus tracking reveals actin-aided viral trafficking of SARS-CoV-2 on the plasma membrane"

**This PDF file includes:**

Materials and Methods

Supplementary Figures 1 to 10

Supplementary Video 1 to 2

**Other Supplementary Materials for this manuscript include the following:**

Supplementary Video 1 to 2

### 1 Experimental Set-up

#### 1.1 Instrumental Set-up Overview

A detailed instrumental diagram is depicted in Supplementary Figure 1 and consists of 3D-SMART tracking excitation optics and 3D-FASTR imaging excitation optics coupled through a commercial confocal microscope (Zeiss LSM 410, modified by LSM Tech), piezoelectric stage and microscope objective to join both setups together. The microscope is controlled by custom LabVIEW code.

##### 1.1.1 Modification in 3D-TrIm beam path

In 3D-FASTR, the axial range is ~ 8 μm, which is determined by the magnification of the system and the working range of ETL. When the 3D-SMART and 3D-FASTR beams are collimated, with the tracking beam defined as the 0 μm axial reference, the ETL scan covers a range extending 6 μm above the particle but only 2 μm below it. This configuration tends to oversample the buffer region while under-sampling the cellular environment beneath the particle. To address this limitation, in previous configuration of 3D-TrIm, the focus was biased to move the tracking volume closer to the center of the imaging volume to more frequently image areas beneath the particle, leading to a slightly converging behavior of the tracking beam. The net impact is with the tracking beam serving as the 0 μm reference, the axial depth range of the ETL scan extends 5 μm below the particle, and only 3 μm above it. This balances the need to sample the cell more aggressively on approach while maintaining 3D cellular imaging when particles are bound or internalized.

In the new optical configuration of 3D-TrIm, the tracking beam was collimated and the imaging volume was shifted downward by replacing several lenses with longer focal lengths, accompanied by corresponding adjustments to the downstream optical components. A negative 500 mm meniscus lens (**NL**. Thorlabs, LF1988-B) was added in series with the ETL to shift the focal plane. To compensate for this focal shift and maintain the ability to focus on PD1, a new 200 mm positive lens (**LPD1**. Thorlabs, LA1708-B) was installed. Additionally, the slider lens was replaced with a weaker 2000 mm lens (**L2**. Thorlabs, LA1258-B) from the original 1000 mm to optimize system performance. To provide a sufficiently large working range for axial scanning, the objective lens was replaced with a model offering the same magnification and numerical aperture (**OL**. Nikon Plan Apo VC 100 ×) but with an extended working distance.

Additionally, a shutter on/off feature was implemented to extend the imaging duration by stopping the two-photon excitation and decreasing the volume rate.

#### 1.2 Sample preparation and validation

For these single-virus tracking experiments, we incorporated fluorescent protein into SARS-CoV-2 D614G VLPs by fusing StayGold to HIV-1 Vpr which is packaged within the pseudovirus nucleocapsid. 3D-TrIm trajectories were acquired on cells labeled with SYTO 61 (targeted to nucleic acids). The cell label was chosen to maximize chromatic separation between the tracking and imaging, and the largest contributor to tracking crosstalk was cell autofluorescence by singlephoton excitation, which was low enough not to perturb the active-feedback single-virus tracking. We cultured monolayers of hACE-negative ***293T/17***, hACE2-expressing ***293T-hACE2-low*** and ***293T-hACE2-high***, and hACE2/TMPRSS2 coexpressing ***293T-ACE2.TMPRSS2*** cells. This selection of cell types offered variety in cell surface receptors/coreceptors and concentrations to observe their influences on the early stages of viral infection.

##### 1.2.1 Production of StayGold Labelled VLPs with SARS-CoV-2 Spike Protein

The pseudotyped lentiviruses with the SARS-CoV-2 spike protein and StayGold tagged Vpr (CoV2-StayGold VLPs) were produced by transfecting the target cells with corresponding lentiviral plasmids as previously reported. Before the production of VLPs, 293T/17 cells were grown in completed RPMI-1640 (Millipore Sigma, no. R8758) supplemented with 10% tetracycline-free FBS (Neuromics, no. FBS002-T) and 1× penicillin–streptomycin (Corning, no. 30-002-CI). The completed RPMI-1640 was also used as the transfection medium and the medium used to collect the virus.

CoV2-StayGold VLPs were produced using Lenti-X Packing Single Shots (D614G Spike, Truncated; Takara Bio, no. 632675), with pLVXS-ZsGreen1-Puro (Takara Bio, no. 632677) as the lentiviral vector plasmid DNA and equal amounts of additional Vpr-StayGold plasmid.

The VLPs produced were concentrated with a Lenti-X Concentrator (Takara Bio, no. 631231) and were resuspended with PBS (Genesee Scientific, no. 25-507). Lentivirus titration was determined by p24 ELISA using a Lenti-X p24 Rapid Titer Kit (Takara Bio, no. 632200) and a plate reader (PerkinElmer Victor3 V). The lentiviral titers were reported in infectious units per mL (IFU mL–1), typically 5–8 × $10^9$ IFU $mL^{-1}$. The VLP stock solutions were stored in aliquots at −80 °C.

##### 1.2.2 Antibody labelling

To produce a labeled secondary antibody for immunofluorescence assays, 50 μg of goat anti-mouse IgG H&L (Abcam, #ab6708) or goat anti-rabbit IgG H&L (Abcam, #ab6702) was dialyzed against 1 L of 1× PBS at 4 °C for 4 h (D-tube mini, MWCO 6-8 kDa, Millipore Sigma, #71504-M). The volume of the recovered antibody was measured and to this 1/10 the volume of 100 mM $NaHCO_3$ (pH = 8.3) was added. Next, to initiate labeling 5-fold molar excess of AF555-NHS ester (ThermoFisher, no. A20009), dissolved and subsequently diluted in anhydrous DMSO, was added so that the final volume of DMSO was < 2%. The reaction was left to continue at room temperature for 1 h. The solution was applied to a desalting column (Zeba™ Spin, 7K MWCO, ThermoFisher, #89882) pre-equilibrated with 10 mM Tris-HCL, pH = 7.5, and 0.01% $NaN_3$, to terminate the reaction and remove uncoupled dye.

##### 1.2.3 Immunofluorescence

The packing efficiency of Vpr.StayGold inside the SARS-CoV-2 pseudotyped VLPs was analyzed by two different immunofluorescence assays, one targeted against the inner capsid the other against the external envelope glycoprotein. VLPs were adhered to autoclaved glass coverslips (VWR, # 48380-046) coated with poly-L-lysine (#P6282, Millipore Sigma), overnight in PBS at 4°C.

VLPs were first fixed for 20 min using 7.4% formalin (Fisher, no. BP531-500) in PBS. Subsequently, washed (3×, 5min) and permeabilized for 10 min in T-PBS (0.1% Triton X-100 in PBS (pH=7.2)). Coverslips were blocked with buffer containing 10% normal goat serum (MP Biomedicals, #IC19135680), 0.2 M Glycine, and 0.1% Triton X-100 in PBS (pH=7.2) for 90 min at room temperature.

VLPs were stained for capsid protein using mouse anti-HIV-1 p24 gag monoclonal antibody (The following reagent was obtained through the NIH HIV Reagent Program, Division of AIDS, NIAID, NIH: Anti-Human Immunodeficiency Virus 1 (HIV-1) p24 Gag Monoclonal (#24-3), ARP-6458, contributed by Dr. Michael Malim), at 2 ng $\mu L^{-1}$ in blocking buffer 120 min at room temperature. Coverslips were washed (3×, 5 min) in T-PBS to remove any unbound primary antibody.

VLPs were stained for SARS-CoV-2 envelope glycoprotein using polyclonal rabbit anti-SARS-CoV-2 spike glycoprotein (BEI Resources, NR-52947), at 1000× dilution in blocking buffer 120 min at room temperature. Coverslips were washed (3×, 5 min) in T-PBS to remove any unbound primary antibody.

As a secondary antibody, AF555-labelled Goat Anti-Mouse (p24)/Rabbit (spike protein) IgG (described above) was used at 4 ng $\mu L^{-1}$ in blocking buffer for 120 min at room temperature. Finally, coverslips were washed with T-PBS (3×, 5 min), and mounted in PBS. In control experiments the same procedure was followed except either the primary or secondary antibody incubation was omitted.

Immuno-stained VLPs were imaged on a spinning disk confocal (Andor Dragonfly 505) on a Leica DMi8 inverted microscope using 100x/1.40-0.70 HCX PL APO (Leica 11506210) oil objective and 488 nm and 561 nm laser lines for excitation (40 µm pinhole). Images were captured on an Andor iXon Life 888 1024×1024 EMCCD camera, and the system was controlled by Fusion 2.0. (Duke University Light Microscopy Core Facility NIH Shared Instrumentation grant 1S10RR027867-01).

Intensity based colocalization was performed using IMARIS software (Oxford Instruments) to extract the Pearson's coefficient for each field of view. Independently, Vpr.StayGold centers were identified in MATLAB (MathWorks) by determining the maximum intensity of each foci and the intensity at the corresponding position in the AF555 channel.

###### 1.2.4 VLP characterization via 3D-SMART

The size and brightness of Vpr.StayGold or Vpr.eGFP incorporated VLPs was evaluated by real-time 3D tracking to extract the diffusion coefficient and particle emission rate. Free virus particles were tracked in HEPES pH = 7.4 buffered solution (live cell imaging solution, ThermoFisher, no. A14291DJ) at room temperature. The tracking microscope configuration was identical to that used for tracking and imaging as previously described. The average excitation power of the 488 nm laser was ~200 nW at the focus.

###### 1.2.5 Production 293T-hACE2 by lentiviral transduction

To generate 293T cells stably expressing hACE2, cells were transduced with the respective lentivirus (pWPI-IRES-Puro-Ak-ACE2, kindly provided by Dr. Sonja Best, Addgene viral prep #154985-LV) at gradient MOI. At 48 hpi, positively transduced cells were selected with puromycin (1 μg/mL) for 5 days and the resistant cells amplified and stored in liquid nitrogen.

##### 1.2.6 Fluorescence-activated cell sorting (FACS)

Following lentiviral transduction of 293T cells with a human ACE2 expression construct, cells were stained with human ACE-2 Alexa Fluor 647-conjugated antibody (R&D Systems, no. FAB9332R). FACS was performed on a Beckman Coulter Astrios cell sorter to isolate the high-ACE2-expressing cells and expanded for downstream experiments. ACE2 expression levels in the sorted population were confirmed by flow cytometry using the same protocol (Supplementary Figure 4a).

##### 1.2.7 Infection assay

To assess the infectivity of SARS-CoV-2 pseudotyped VLPs containing fluorescent Vpr, different 297T-derived cells were inoculated with SARS-CoV-2 lentiviral vector encoding the ZsGreen1 reporter construct pLVXS-ZsGreen1-Puro. 297T-derived cells were grown using complete DMEM which comprised of DMEM Media (Corning, no. 10-013-CV) supplemented with 10% FBS (Millipore Sigma, no. F2442), and 1 × penicillin-streptomycin (Corning, #30-002-CI) in T12.5 flask (Genesee Scientific, no. 25-205). Cells were plated in complete DMEM at $5\times10^4$ cells/well in an 8 well μ-slide, glass bottom (Ibidi, no. 80827). 297T/17 cells were immediately inoculated with a multiplicity of infection (MOI) of 1, 10, or 100 transfecting units per cell of lentiviral vector. The infected cultures were incubated at 37 °C in an atmosphere of 5% $CO_2$.

At 72 h p.i., cells were fixed with 4% paraformaldehyde (PFA) solution in 1 × PBS (Ph = 7.2) for at least 20 minutes. Then the cells were stained with 1.5 μM of Hoechst 34580 (ThermoFisher, no. H21486) diluted with 1 × PBS. The fixed sample was kept at 4 °C until imaging within 3 days.

The imaging was conducted on a spinning disk confocal microscope (Andor Dragonfly 505), equipped with 40×/1.3 HC PL APO CS2 (Leica 11506358) oil objective, and 405 nm and 488 nm laser lines for excitation (40 μm pinhole). Simultaneous two-color imaging was performed at 2.5 % and 0.5 % 405 nm and 488 nm laser power, respectively. Images were captured on an Andor iXon Life 888 EMCCD camera with 200 msec exposure time for both colors.

#### 2 Analytical Methods

##### 2.1 Mean-square displacement analysis for particle diffusion

The diffusion coefficient can be obtained by linear fitting of the mean square displacement (MSD) with lag time (τ). The MSD was calculated using the definition:

$$MSD(\tau) = N^{-1}\Sigma_{n=1}^{N}\left((x(n+\tau)-x(n))^2) + (y(n+\tau)-y(n))^2) + (z(n+\tau)-z(n))^2)\right) \quad (2)$$

Here, $x(n)$, $y(n)$, and $z(n)$ are the coordinates of the trajectory at timepoint $n$. $N$ is the total number of data points, which corresponds to the window size in sliding window analysis.

Trajectories were quantified using diffusion coefficient ($D$), power exponent α, and packing coefficient ($Pc$). A power-law MSD analysis is typically applied for anomalous diffusion:

$$MSD(\tau) = 2nD\tau^{\alpha} \tag{3}$$

where n is the number of dimensions, D is the diffusion coefficient, τ is the lag time, and the exponent α is used to classify anomalous diffusion (α < 1: subdiffusion; α = 1: Brownian motion; α > 1: superdiffusion).

The hydrodynamic radius (r) of the particle being tracked was calculated using the Stokes-Einstein relation:

$$D = \frac{k_B T}{6\pi\eta r} \tag{4}$$

Here, $k_B$ is the Boltzmann constant, $T$ is temperature, and η is viscosity of solution.

For drift velocity calculations of linear trafficking events, MSD was fitted with a different model:

$$MSD(\tau) = 2nD\tau + v^2\tau^2 \tag{5}$$

where $v$ is the drift velocity, which describes a net displacement or 'drift' over longer time scales. The minimum trustworthy drift velocity calculated from this algorithm is 0.4 nm/sec, which corresponds to 40 nm displacement over 100 seconds and is determined by the tracking localization precision of 3D-TrIm.

#### 2.2 Cell Boundary Determination

Image segmentation of the cell boundaries was required for calculating virus-cell distances. Membrane stains were considered as a method for identifying the edge of the cell, but the continual recycling of the membrane by the live cells led to a complete internalization of any membrane dye within 30 minutes of the start of the experiment. To avoid perturbing actin dynamics, actin staining was not performed, as the goal of the experiment was to examine the role of actin activity in viral membrane trafficking. Therefore, SYTO (which labels the cytosol and nucleus through DNA/RNA staining) was used in this work and it was stable and reliable indicators of cell morphology.

To segment the cell boundaries in the image, we utilized the morphological top-hat filter with a spherical structuring element of radius 5 pixels using the `imtophat` function in MATLAB (MathWorks) and applying this transformation to the local volume intensity. This function retains elements smaller than the structuring element and brighter than the background. To remove segmented noise pixels, we employed a second morphological filter, `bwareaopen` in MATLAB, which removes small unconnected objects. Finally, the segmented volume is passed to the `isosurface` function in MATLAB to determine the faces and vertices of the resulting segmentation.

2.3 Cell-to-Virus Distance Calculation

To determine the minimum distance, *d*, between each point, *i*, in the cellular image, *c*, and the viral coordinates, *v*, the volume must be segmented using the top hat procedure described above. Then, for each point ($i$) of the trajectory, the closest voxel coordinate of cellular surface ($i\_min$) was identified using MATLAB functions `KDTreeSearcher` and `knnsearch`. The minimum Euclidean distance shown in the formula below is taken as the distance between the viral particle to the cell surface:

$$d_i = \sqrt{\left(c_{x,i\_min} - v_{x,i}\right)^2 + \left(c_{y,i\_min} - v_{y,i}\right)^2 + \left(c_{z,i\_min} - v_{z,i}\right)^2} \qquad (6)$$

To avoid the artifacts induced by the sample drift or cellular morphological change over the long tracking duration, for each time points, the distance calculation was performed on the average of three closest volumes.

Surface events were defined using the calculated virus-to-surface distances between ± 1 μm, on the doubled size scale as the imaging Z-PSF.

2.4 Simulation of a 3D random walk along a cylindrical surface

A 3D random walk trajectory was simulated along the surface of a cylinder aligned parallel to the X-axis. At each time step, the particle underwent a random displacement along the X-axis and a random angular displacement around the cylindrical surface. The corresponding Y and Z positions were calculated based on the updated angular position and a constant cylinder radius. In total 160,000 data points were simulated for the trajectory to mimic the duration and number of localizations of the experimental data. To approximate experimental measurement conditions, Gaussian-distributed localization noise with standard deviations of 20 nm, 20 nm, and 80 nm was added to the X, Y, and Z coordinates (determined by experimental localization precision), respectively. The simulated trajectory is shown in Supplementary Figure 6a.

A 2D heatmap is generated for the YZ view of the simulated results, in which two bright spots appear on the side while the middle is blurred out due to worse precision (Supplementary Figure 6b). By performing PCA on experimental trajectory in Figure 2, a heatmap of PC2 and PC3 view was compared to be similar to the YZ view of simulated result (Supplementary Figure 6c).

For the 3D point cloud color-coded by density shown in Figure 2, it is done by using a radius-based range search using a search radius of 50 nm. For each point, the number of neighboring points within this radius was counted to estimate local point density.

2.5 Criteria for resolving state transitions

Different states were determined using a combination of the particle diffusivity trace (D), distance trace (Dist.), drift velocity trace (V), particle intensity trace (I), diffusion pattern, and image intensity. Detailed criteria for state categorization are listed below:

| State | Diffusion coefficient | Drift Velocity | Distance |
|---|---|---|---|
| Free diffusion | $D \geq 1\ \mu m^2/s$ | $V < 0.4$ nm/s | Any distance |
| Bound to cell | $D_{bound} < 10 \times D_{free}$ | $V < 0.4$ nm/s | Dist. < 1 μm |
| Membrane trafficking | $1.6 \times 10^{-4} < D \leq 0.05$<br>Linear motion | $V \geq 0.4$ nm/s | -1 μm < Dist. < 1 μm |
| Internalized trafficking | $1.6 \times 10^{-4} < D \leq 0.05$<br>Linear motion | $V \geq 0.4$ nm/s | Dist. ≤ -1 μm |

2.6 Volume Render Visualization

3D visualization of intensity and distance volume data was performed using Amira 3D 2021.1. Binarized arrays with registered spatial coordinate data were imported into the program and visualized using cubic interpolation and alpha composition. Alpha mapping was controlled through a gamma curve to reduce the opacity of noisy features. Features were rendered using an opacity threshold value which solidifies only surface regions and not background, typically by using a value of ~ 0.2. Trajectories were imported as comma-delimited spreadsheets containing coordinate, time, diffusivity, drift velocity, and distance data and displayed as Line Set view objects with a line width of 2.

#### 3 Supplementary Figures

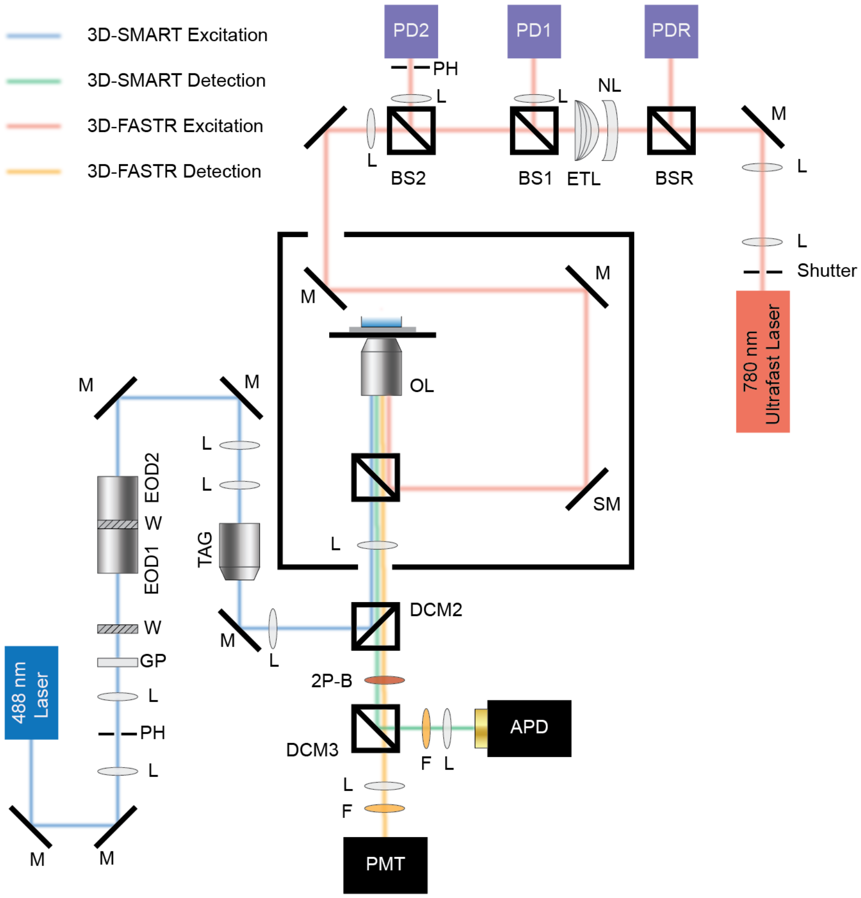

**Supplementary Figure 1: Instrument diagram**

M: mirror, L: lens, PH: pinhole, GP: Glan-Thompson Polarizer, W: half wave plate, EOD: electro-optic deflector, TAG: tunable acoustic gradient lens, DCM: dichromatic mirror, OL: objective lens, F: fluorescence emission filter, BS: beam splitter, ETL: electrical tunable lens, NL: negative lens, PD: photodiode, 2P-B: two-photon blocker, SM: scanning mirror, APD: avalanche photodiode.

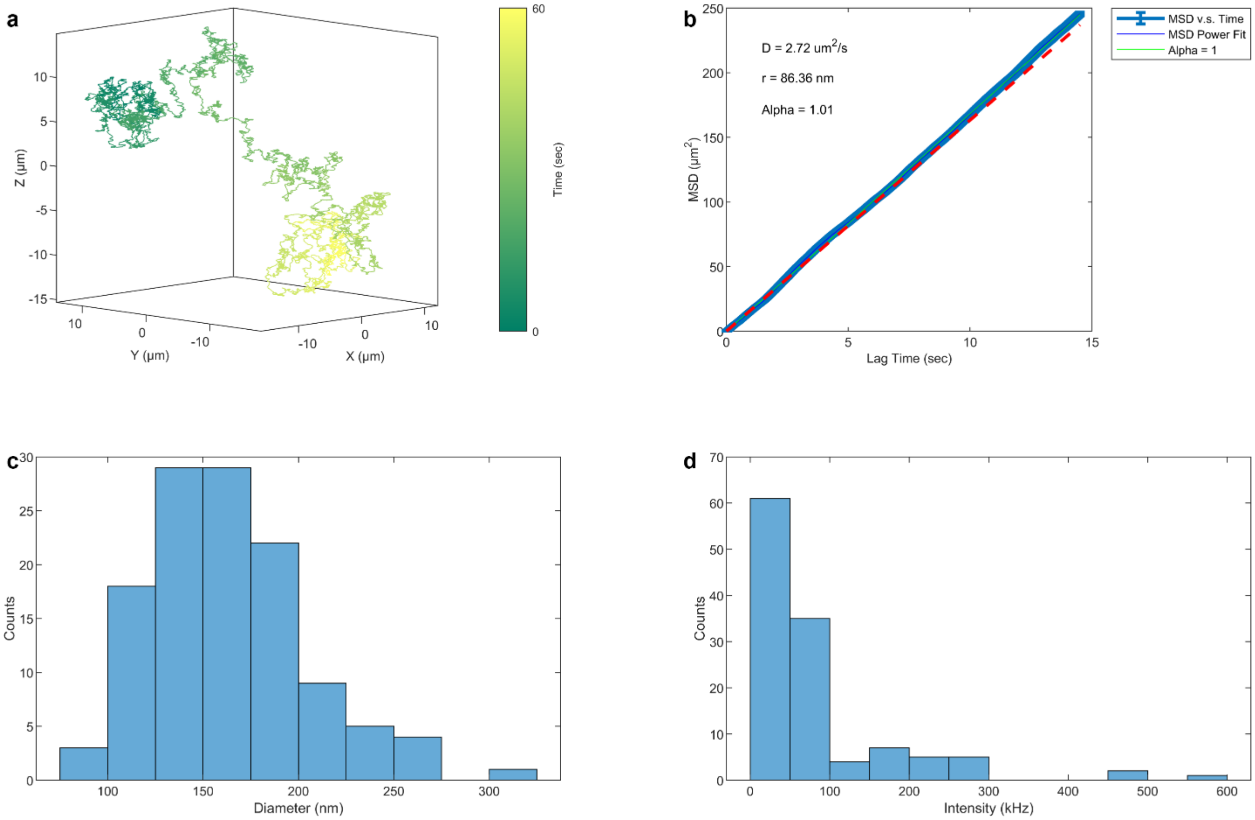

**Supplementary Figure 2: Characterization of CoV2/SG-VLPs using SVT without cells**

(a) An example trajectory of a free diffusing CoV2/SG-VLP in buffer.
(b) MSD analysis of the trajectory shown in (a).
(c) Hydrodynamic radius distribution of CoV2/SG-VLPs.
(d) Intensity distribution of CoV2/SG-VLPs. 162 ± 41 nm (mean ± std), N = 120.

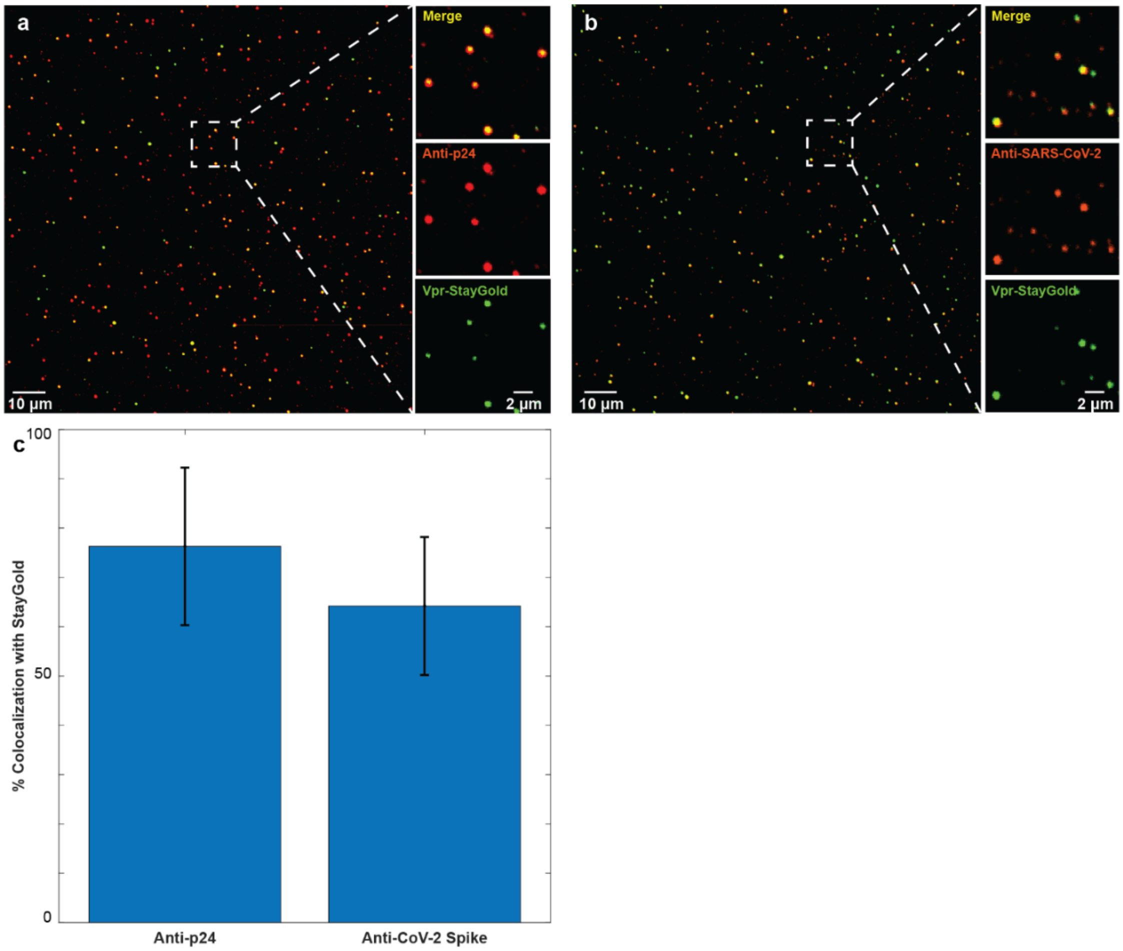

**Supplementary Figure 3: Successful incorporation of Vpr-StayGold into VLPs with SARS-CoV-2 spike protein validated by immunofluorescence**

(a) Representative image of anti-p24 immunofluorescence of CoV2/SG-VLPs incubated with primary and secondary antibodies. False-color merge of Vpr-StayGold (green) and AF555-labelled p24 (red). A zoom-in view is shown in the right panel: Top: merge of two channels below. Middle: channel of AF555-labelled p24. Bottom: channel of Vpr-StayGold.

(b) Representative image of anti-CoV-2-spike immunofluorescence of CoV2/SG-VLPs incubated with primary and secondary antibodies. False-color merge of Vpr-StayGold (green) and CF555-labelled CoV-2-spike protein (red). A zoom-in view is shown in the right panel: Top: merge of two channels below. Middle: channel of CF555-labelled CoV-2-spike protein. Bottom: channel of Vpr-StayGold.

(c) The colocalization efficiency of anti-p24 and anti-VSV-G with StayGold signal is 76.3% ± 16.0% and 64.2% ± 14.0% (mean ± std, N = 3), respectively. The colocalization efficiency was calculated by dividing the number of particles with colocalized red and green signals in (a) and (b) by the sum of colocalized and green-signal only particles (corresponding to the total amount of 'trackable' particles).

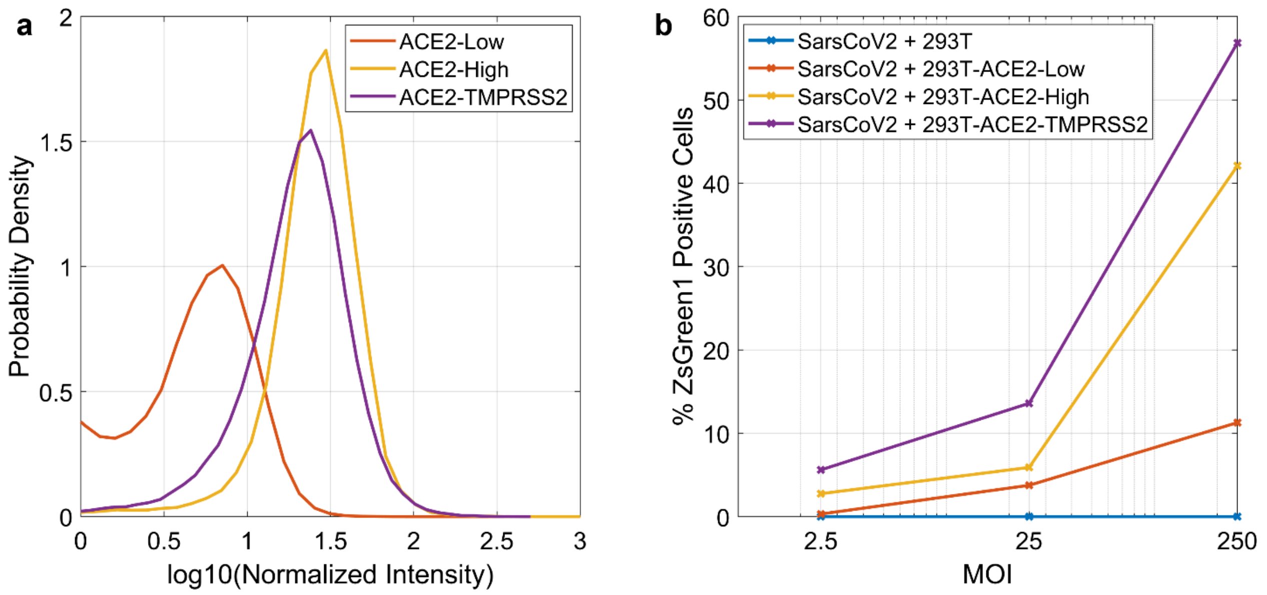

**Supplementary Figure 4: Infection assay results across different cell lines**

(a) Expression level of ACE2 measured on different cell lines through flow analysis. Normalized intensities were calculated by scaling relative to ACE2-negative 293T cells as $I_{norm} = I_{Pos} \div I_{Neg}$. (b) Infection assay performed on different cell lines. These results also indicate a different expression level of hACE2 in 293T-ACE2-low (produced in house by lentiviral transduction) and 293T-ACE2-high (from BEI Resource).

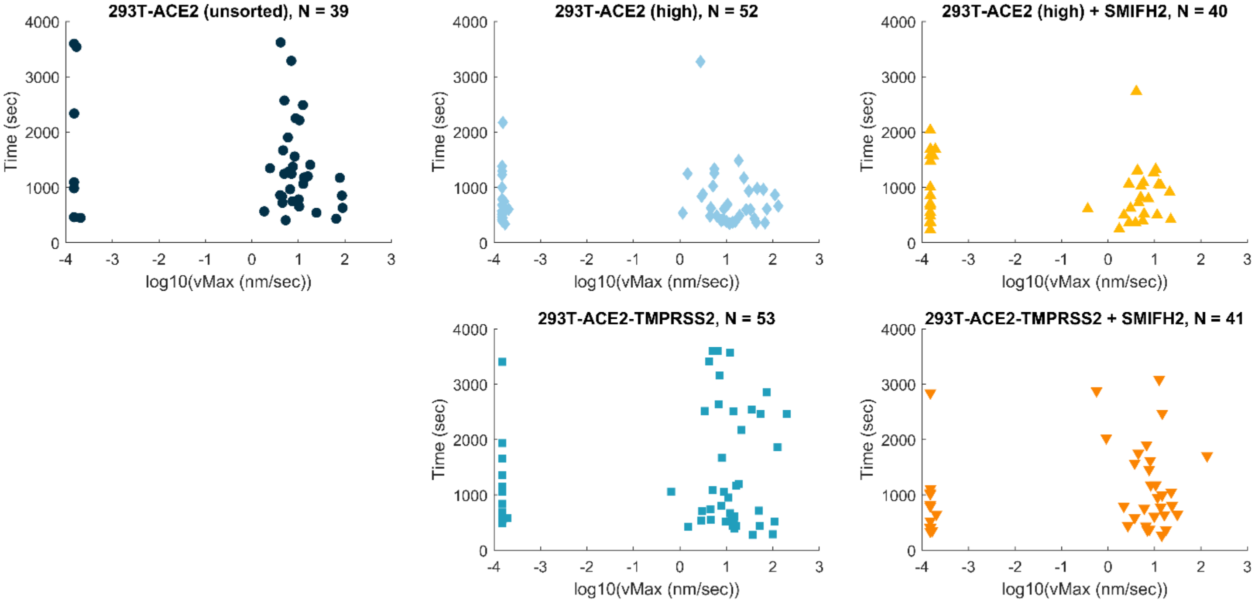

**Supplementary Figure 5: Tracking duration and maximum drift velocity for different experimental conditions**

Scatter plots of log maximum drift velocity (nm/sec) and tracking duration for each experimental condition. The population of log maximum drift velocity close to -4 represents immobilization.

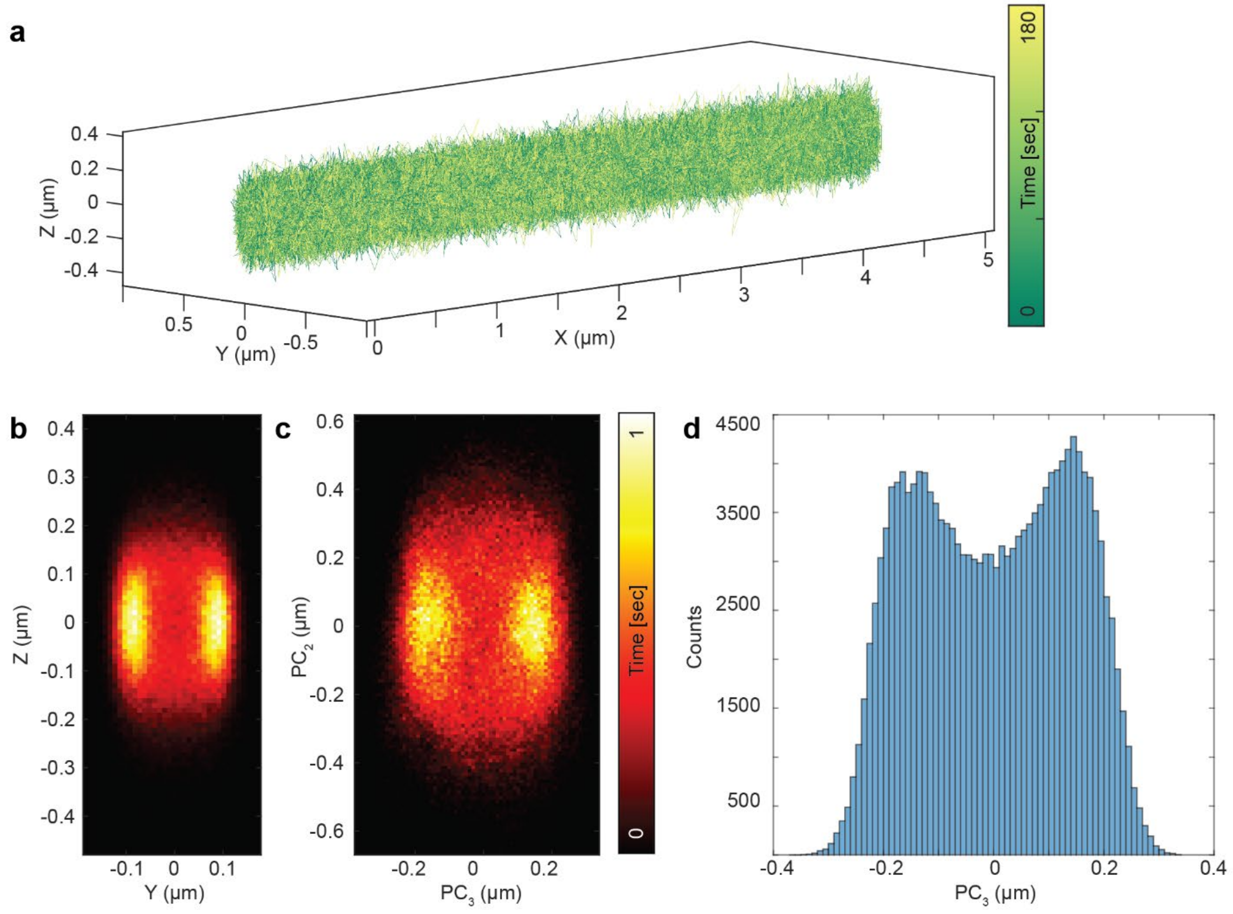

**Supplementary Figure 6: Validation of 3D virus motion on protrusions, related to Figure 2**

(a) Simulated random walker on a 3D cylindrical surface along X axis.
(b) 2D heatmap (YZ view) of the simulated trajectory.
(c) 2D heatmap (PC2 and PC3 view) of the experimental data shown in Figure 2.
(d) Histogram of localization distribution of PC3.

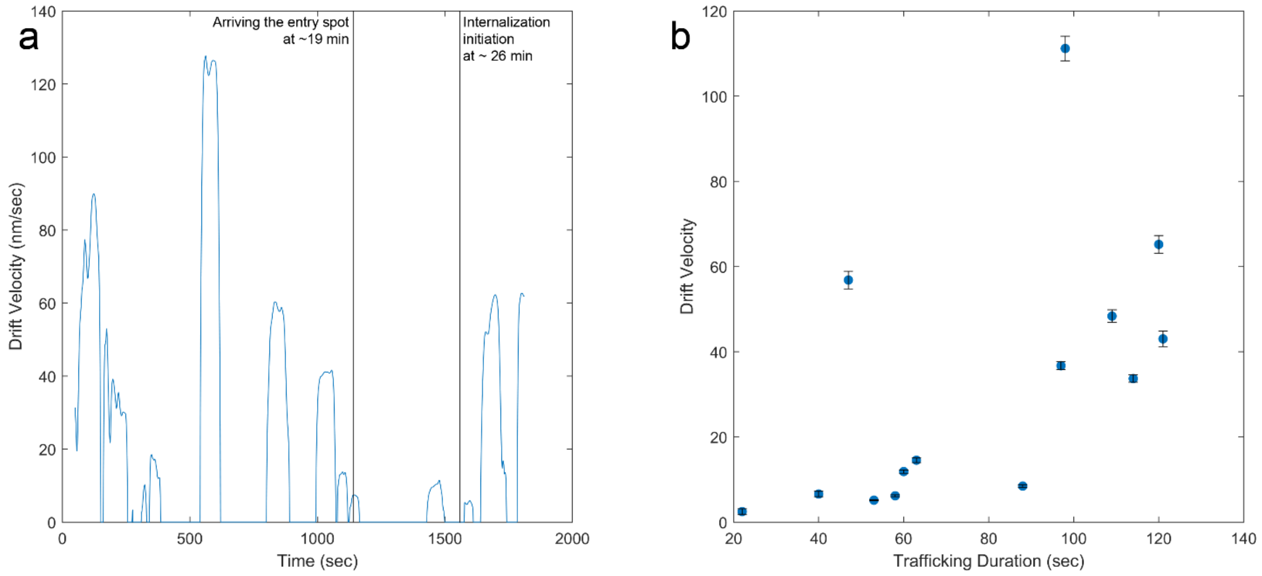

**Supplementary Figure 7: Drift velocities and trafficking durations of membrane trafficking event, related to Figure 3**

(a) Calculated drift velocity shows stop-and-go pattern for membrane trafficking events. The calculation is performed in 100-sec windows and 1-sec steps.

(b) Scatter plot of trafficking duration and drift velocity. Each dot represents the mean of drift velocity throughout each trafficking event, and the error bar represents the S.E.M.

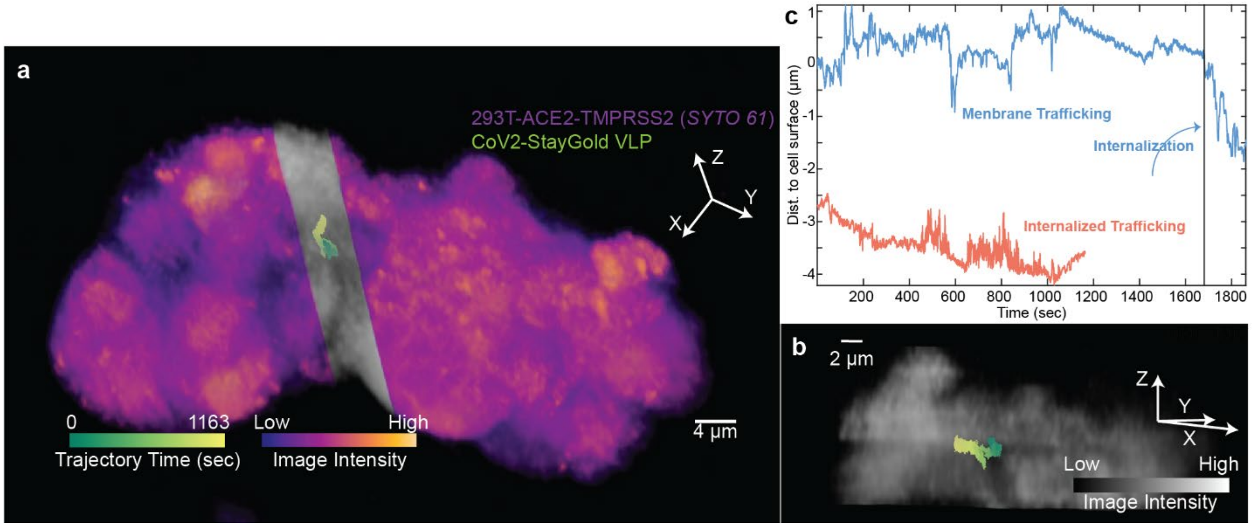

**Supplementary Figure 8: A comparison between membrane trafficking and internalized trafficking, related to Figure 3**

(a) An example of internalized VLP trafficking. Two slice planes were used to section the YZ plane and create a channel that renders the interior volume section semitransparent, revealing the trajectory embedded beneath the cell surface.

(b) Side view demonstrates that the VLP is internalized deep within the cell.

(c) A comparison of the virus-to-cell distance traces for membrane trafficking followed by internalization (Figure 3A-C, blue) versus internalized trafficking (a-b). Distance traces are a 1-sec running mean.

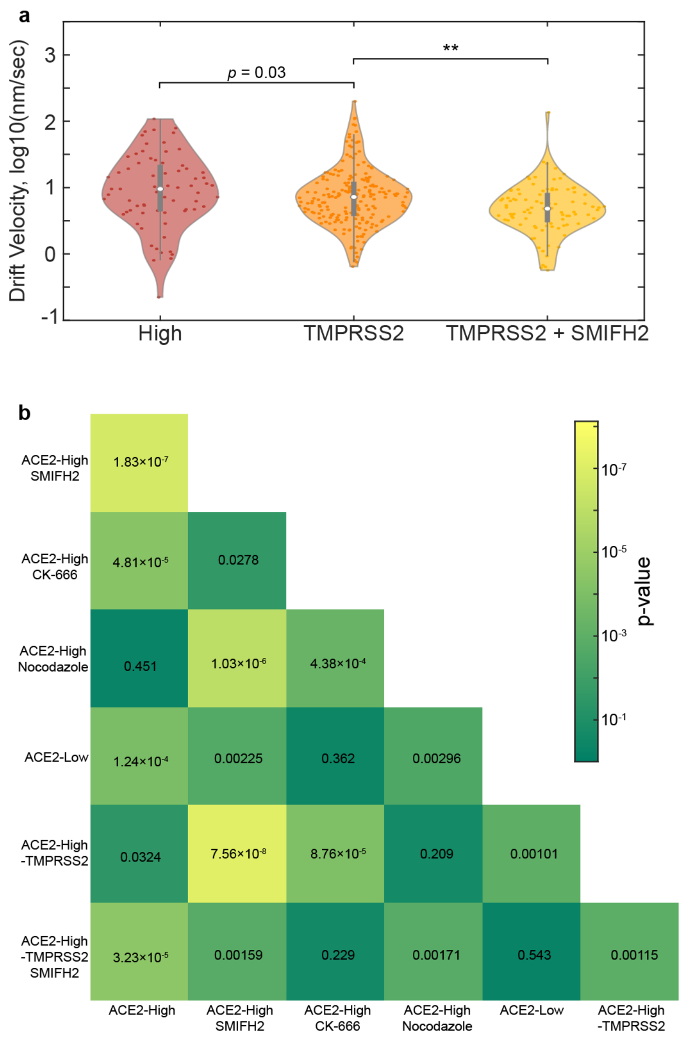

**Supplementary Figure 9: 293T-ACE2-TMPRSS2 cells are less sensitive to SMIFH2 treatment**

(a) Violin plots of drift velocity, expressed as $\log_{10}$ (nm/sec), of CoV2/SG-VLPs in comparison of the existence of TMPRSS2 and SMIFH2 treatment. Drift velocities are calculated in 100-second segments and immobilized segments are removed from this analysis. $N = 70, 205, 93$, respectively. **: $p<0.01$, Kolmogorov-Smirnov test.

(b) Heatmap of pairwise p-values for the drift velocity comparisons across different conditions.

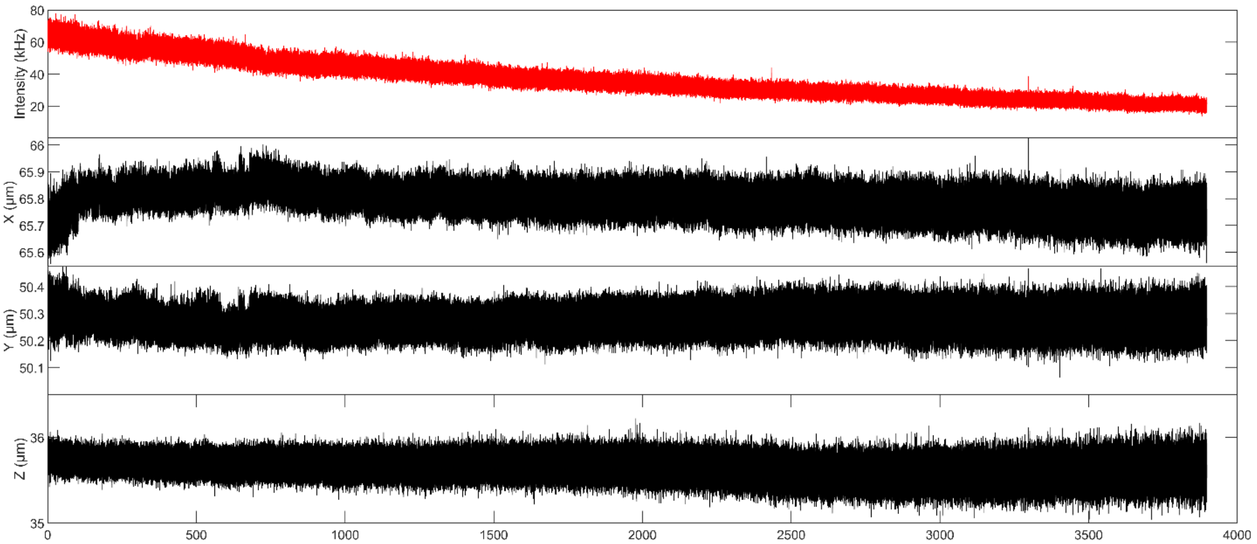

**Supplementary Figure 10: Minimal sample drift observed during long 3D-TrIm experiments**

(a) A ~ 3900-sec trajectory fixed on the coverslip indicates minimal sample drift during hour-long 3D-TrIm experiments.

(b-d) Intensity, X, Y, and Z trace of this trajectory.

#### 4 Supplementary Movie Captions

**Supplementary Video 1: An eventful single SARS-CoV-2 VLP tracking trajectory, related to Figure 2.**

3D reconstruction of live 293T cells with high expression level of ACE2 receptors (stained with nucleic acid label SYTO61) co-rendered with virus trajectory. Trajectory (~ 887 sec) is segmented into 25 segments per second (25 frames per second when playback rate is 1×) and color mapped by time. The progress bar shows how the trajectory is further categorized: (1) Free diffusion period (playback rate: 1×); (2) Viral surfing on cellular protrusion (playback rate: 10×); (3) Membrane trafficking (playback rate: 40×). The end points of trajectories segments are labelled with spheres (refreshing rate is consistent with the trajectory, i.e., 25 fps at 1× playback rate), representing the position of the viral particle. Image volumes formed from maximum intensity projection over time from local volumes acquired over 16 frame-times. Cells are color-coded by image intensity.

**Supplementary Video 2: SARS-CoV-2 VLP actively trafficked on cellular membrane, related to Figure 3.**

3D reconstruction of live 293T cells with high expression level of ACE2 receptors and TMPRSS2 co-receptors (stained with nucleic acid label SYTO61) co-rendered with virus trajectory. Trajectory (~ 1862 sec) is segmented into 25 segments per second (25 frames per second when playback rate is 1×) and color mapped by the distance between virus and cell surface (green: distance > 0, white: distance = 0, blue: distance < 0). The playback rate is constant 30×. Image volumes formed from maximum intensity projection over time from local volumes acquired over 16 frame-times. Cells are color-coded by image intensity. We observed the VLP undergoing trafficking on cellular membrane and eventually was internalized.
